# The presence and induction of regioselective dehydroxylases dictate urolithin metabolism by *Enterocloster* species

**DOI:** 10.1101/2025.08.06.668933

**Authors:** Reilly Pidgeon, Arianna Giurleo, Lharbi Dridi, Bastien Castagner

## Abstract

Urolithins are a class of bioactive metabolites derived from the metabolism of dietary ellagitannins by the human gut microbiota. In the gut, urolithins are dehydroxylated regioselectively based on microbiota composition and activity. A single 9-hydroxy urolithin dehydroxylase (*ucd*) operon in gut resident *Enterocloster* species has been described to date; however, most enzymes in the urolithin metabolic pathway remain uncharacterized. Here, we investigate urolithin cross-feeding between members of the gut microbiota and discover a novel urolithin dehydroxylase in a subset of *Enterocloster* species. We show that urolithin intermediates, released by gut resident *Gordonibacter* species during ellagic acid metabolism, are dehydroxylated at both the 9- and 10-positions by *E. asparagiformis*, *E. citroniae*, and *E. pacaense*, but not *E. bolteae*. Using untargeted proteomics, we uncover a 10-hydroxy urolithin dehydroxylase operon, termed *uxd*, responsible for these species-specific differences in urolithin metabolism. By inducing *uxd* expression with diverse urolithins, we show that 9-hydroxy urolithins are required for *uxd* transcription and 10-position dehydroxylation. Collectively, this study reveals some of the genes, proteins, and substrate features underlying differences in urolithin metabolism by the human gut microbiota.

## Introduction

The gut microbiota contributes to host homeostasis by regulating immune function, intestinal barrier integrity, and nutrient availability ^1^. Due to the immense diversity of biosynthetic and metabolic gene clusters present in gut bacteria ^2^, many ingested xenobiotics (food, drugs, pollutants) are metabolized by bacterial enzymes through reactions that the host alone cannot perform ^3–5^. Urolithins (*6H*-dibenzo[*b*,*d*]pyran-6-ones) are a class of postbiotics derived from the metabolism of dietary ellagitannins by gut bacteria. The most common urolithin metabolite detected in host biological fluids following ellagitannin consumption is urolithin A (3,8-dihydroxy-urolithin, uroA) ^6,7^. In the host, uroA can activate mitophagy to improve mitochondrial health and exercise capacity ^8–10^. This mitophagy-promoting activity triggers immune responses in the context adoptive T cell transfer by expanding T memory stem cells, a subset of minimally differentiated T cells that mediate potent anti-tumor responses ^11^. UroA is also proposed to enhance gut barrier integrity by signaling through the aryl hydrocarbon receptor (AhR) pathway ^12^ and has been shown to reduce *Clostridioides difficile* pathogenesis in murine models ^13^. Based on the abovementioned activities, uroA is currently being evaluated in the clinic for indications in obesity (NCT05921266) ^14^, cancer (NCT06022822), and aging (NCT06619457 and NCT06556706).

Dietary ellagitannins, the large and poorly absorbed precursors of urolithins, are plant secondary metabolites often consumed via berries, nuts, pomegranates, and aged alcoholic beverages ^15^. This family of compounds is characterized by a central glucose (open or closed ring) linked to hexahydroxydiphenoyl (HHDP) and derivative groups. In the gut, the HHDP unit of ellagitannins is hydrolyzed and undergoes spontaneous lactonization, producing ellagic acid (EA) ^16^. EA is decarboxylated and sequentially dehydroxylated by gut bacteria into diverse urolithins, depending on microbiota composition and function ^17,18^. These individual differences in urolithin metabolism have been classified into 3 distinct metabotypes characterized by the terminal metabolites detected in host fluids and feces: 0 (no terminal metabolites), A (only uroA), and B (isouroA, uroB, and can include uroA) ^7^. Like uroA, dietary sources of ellagitannins are in clinical trials for a variety of conditions; therefore, understanding the metabolism of this broad family of compounds is important since treatment responses may vary depending on urolithin metabotypes.

Although urolithin metabolism is prevalent in human populations ^6,19^, few urolithin-metabolizing species have been described in the literature. The Eggerthellaceae family members *Gordonibacter* spp. and *Ellagibacter isourolithinifaciens* convert EA into urolithin M5 (3,4,8,9,10-pentahydroxy-urolithin, uroM5), urolithin M6 (3,8,9,10-tetrahydroxy-urolithin, uroM6), and urolithin C (3,8,9-trihydroxy-urolithin, uroC) via an EA decarboxylase, and 4- and 10-position dehydroxylases, respectively (**Fig. 1a**) ^20,21^. *Ellagibacter isourolithinifaciens* can further dehydroxylate uroC via an additional 8-position urolithin dehydroxylase, yielding isourolithin A (3,9-dihydroxy-urolithin, isouroA) ^22^. A subset of gut resident *Lachnospiraceae* belonging to the *Enterocloster* genus (*E. asparagiformis*, *E. bolteae*, *E. citroniae*, and *E. pacaense*) can metabolize uroC into the bioactive terminal metabolite uroA through a 9-position <u>u</u>rolithin <u>C d</u>ehydroxylase encoded by the *ucd* operon ^23^. We previously demonstrated that *ucd*-encoded proteins (UcdC, UcdF, and UcdO) belong to the xanthine dehydrogenase family and use NAD(P)H as an electron donor to dehydroxylate uroC and related 9-hydroxy urolithins, regardless of the presence or position of other hydroxyl groups on the urolithin scaffold. Intriguingly, a subset of uroC-metabolizing *Enterocloster* spp. can metabolize uroM6 into uroA by dehydroxylating both the 9- and 10-positions (**Fig. 1a**) ^24^. The substrate specificity of the Ucd enzyme complex suggests that another gene cluster encodes a 10-position specific urolithin dehydroxylase in *Enterocloster* spp. Here, we report the discovery of an operon, termed the <u>u</u>rolithin 10-position (<u>X</u>) <u>d</u>ehydroxylase (*uxd*), that encodes xanthine dehydrogenase family proteins responsible for urolithin dehydroxylation at the 10-position. We show that uroM6 is released by EA-metabolizing *Gordonibacter* spp., rendering this substrate available for metabolism by *Enterocloster* spp. By integrating untargeted proteomics, comparative genomics, and targeted gene expression analyses, we identify genes and proteins distinct from *ucd* that enable uroM6 dehydroxylation at the 10-position. Curiously, *ucd* and *uxd* operon expression is co-regulated, requiring a 9-hydroxy urolithin substrate for both 9- and 10-position dehydroxylation. Overall, this study uncovers an additional operon that regioselectively dehydroxylates urolithins and highlights how the regiochemistry of urolithins dictates dehydroxylase operon expression in *Enterocloster* spp.

**Figure 1.**
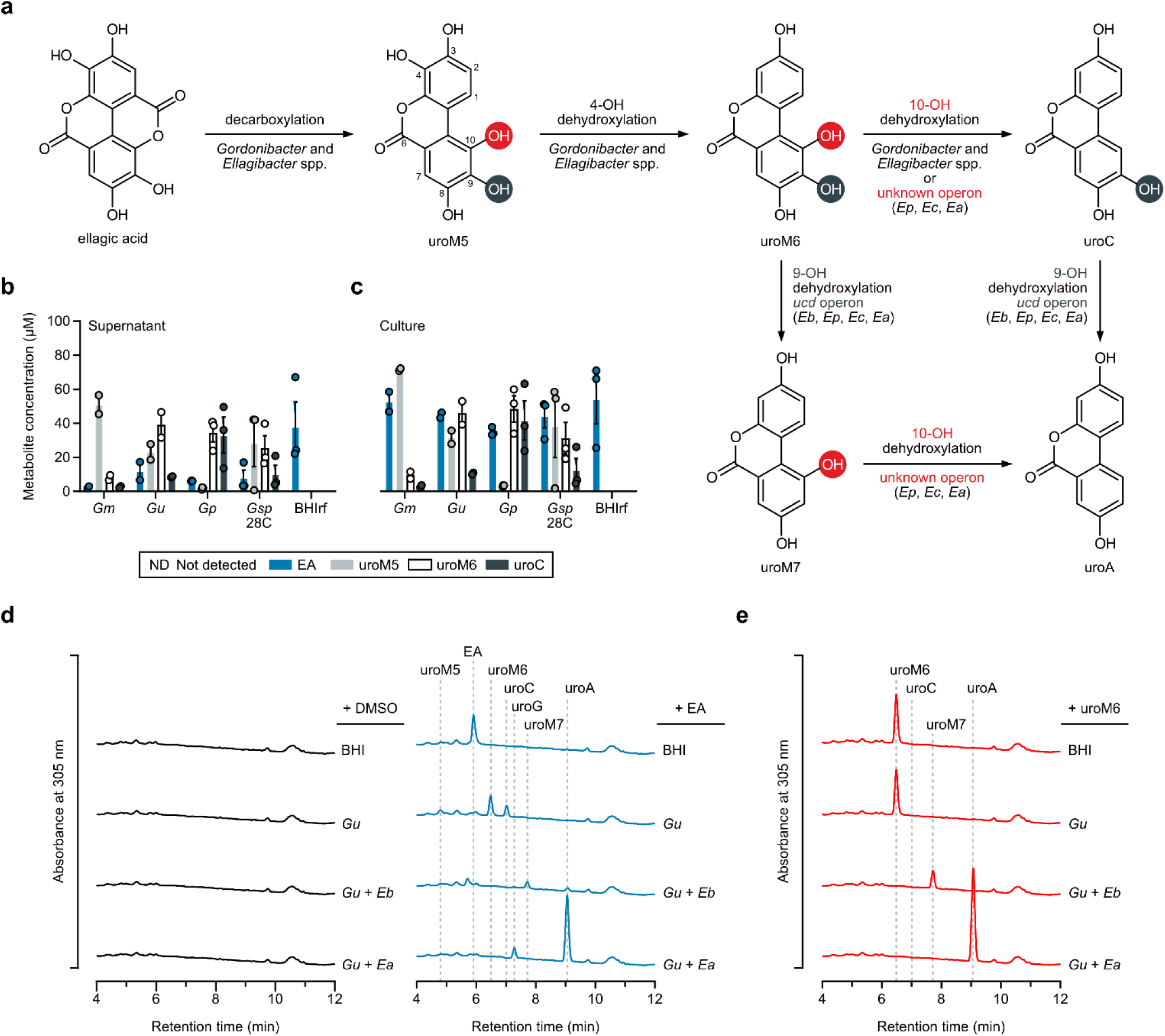
Urolithin intermediates are released by *Gordonibacter* spp. during ellagic acid metabolism. **a** Ellagic acid metabolism scheme. 9- and 10-position urolithin hydroxyl groups are highlighted in blue-grey and red circles, respectively. **b** Quantification of ellagic acid (EA) and urolithin concentrations in cell-free supernatants of *G. massiliensis* (*Gm*), *G. urolithinfaciens* (*Gu*), *G. pamelaeae* (*Gp*), or *Gordonibacter* sp. 28C (*Gsp* 28C) treated with EA (100 μM) (n = 2-3 biological replicates) in BHIrf media after 48 h. Data are represented as means ± SEM. **c** Quantification of ellagic acid (EA) and urolithin concentrations in total cultures of *Gordonibacter* spp. treated with EA (100 μM) (n = 2-3 biological replicates) in BHIrf media after 48 h. Data are represented as means ± SEM. **d** Aligned chromatograms (λ = 305 nm) of urolithins in DMSO (black lines) or EA-treated (100 μM) (blue lines) cultures of *Gu*, *E. bolteae* (*Eb*), and *E. asparagiformis* (*Ea*) in BHI media after 5 days. Retention times of urolithin standards are shown with grey dotted lines. Representative chromatograms from n = 1 of 3 biological replicates. **e** Aligned chromatograms (λ = 305 nm) of urolithins in uroM6-treated (100 μM) (red lines) cultures of *Gu*, *Eb*, and *Ea* in BHI media after 5 days. Retention times of urolithin standards are shown with grey dotted lines. Representative chromatograms from n = 1 of 3 biological replicates.

## Results

### Urolithin M6 is released by *Gordonibacter* spp. and further metabolized by *Enterocloster* spp

Members of the *Gordonibacter* genus convert EA to intermediate urolithins uroM5, uroM6, and uroC (**Fig. 1a**) ^21^. To determine whether 10-hydroxy urolithins (uroM5 and uroM6) were released by *Gordonibacter* spp. during EA metabolism, we incubated *Gordonibacter pamelaeae*, *Gordonibacter massiliensis*, *Gordonibacter sp.* 28C and *Gordonibacter urolithinfaciens* with EA and monitored urolithin production in both whole cultures and supernatants by liquid chromatography-mass spectrometry (LC-MS). The products of EA metabolism were detected in cell-free supernatants after 2 days (**Fig. 1b**), albeit at lower concentrations than those observed in whole cultures (**Fig. 1c**). These data demonstrate that urolithin intermediates derived from EA metabolism are released by *Gordonibacter* spp. And become available to other urolithin-metabolizing bacteria.

To support this hypothesis, we performed co-culture experiments by incubating combinations of *G. urolithinfaciens* and *Enterocloster* spp. (*E. bolteae* or *E. asparagiformis*) with EA or uroM6. Based on the EA decarboxylase and urolithin dehydroxylase activities (4-, 9-, and 10-positions) of these 2-membered communities, we hypothesized that uroA would be detected in co-cultures. As expected, *G. urolithinfaciens* alone metabolized EA to uroM5, uroM6, and uroC (**Fig. 1d**); however, no uroM6 conversion was observed when supplied directly (**Fig. 1e**). In co-cultures of *G. urolithinfaciens* and *E. bolteae* (9-position dehydroxylation only) treated with EA, we could detect the intermediate urolithin M7 (3,8,10-trihydroxy-urolithin, uroM7) and the terminal metabolite uroA (**Fig. 1d**). Direct supplementation with uroM6 yielded uroM7 but no uroA (**Fig. 1e**), demonstrating that only *E. bolteae* can dehydroxylate uroM6 under these conditions. In contrast, co-cultures of *G. urolithinfaciens* and *E. asparagiformis* (9- and 10-position dehydroxylation) treated with EA yielded uroA as a major product and a trihydroxy-urolithin metabolite (**Fig. 1d**). Since we did not have a standard for this metabolite, it was tentatively assigned as urolithin G (3,4,8-trihydroxy-urolithin, uroG), likely derived from the 9- and 10-position dehydroxylation of uroM5 released by *G. urolithinfaciens*. Direct supplementation with uroM6 yielded exclusively uroA (**Fig. 1e**), highlighting differences between activities of urolithin-metabolizing *Enterocloster* spp. Collectively, these data show that simple 2-membered communities can metabolize EA to different urolithins based on dehydroxylase activity and that urolithin features (hydroxylation status and regiochemistry) dictate whether metabolism will occur in whole cells, likely via substrate-specific transporters or dehydroxylase gene induction by urolithin intermediates.

### A distinct dehydroxylase metabolizes 10-hydroxy urolithins

Based on the observation that urolithin intermediates are released by EA-metabolizing *Gordonibacter* spp. and that a subset of *Enterocloster* spp. can metabolize uroM6 to uroA, we sought to identify the genes and enzymes responsible for 10-position urolithin dehydroxylation, as previously done for the 9-position ^23^. We reasoned that either the *ucd*-encoded UcdO (substrate-binding oxidoreductase) subunit of 9- and 10-position-metabolizing species like *E. asparagiformis* was promiscuous, acting at both the 9- and 10-positions of uroM6, or that a separate dehydroxylase was metabolizing the 10-position in these bacteria. Multiple sequence alignment of UcdO proteins from *Enterocloster* spp. (*E. asparagiformis*, *E. citroniae*, and *E. pacaense*) revealed that these sequences were >96.32% identical to the previously characterized *E. bolteae* UcdO (similarities >98.73%) (**Supplementary Fig. 1**). Additionally, predicted urolithin binding site residues W345, Y375, F458, F464, Y536, Y624, and Y632 were conserved among all species (**Supplementary Fig. 1**), suggesting that substrate specificity should be identical in these bacteria. To validate that the *ucd* from *E. asparagiformis* encoded a 9-position-specific dehydroxylase, as observed in *E. bolteae* ^23^, we cloned and heterologously expressed the *E. asparagiformis ucd* operon in *Rhodococcus erythropolis* using a thiostrepton-inducible plasmid ^25^ (pTipQC2-*Ea_ucdCFO*) (**Fig. 2a**). In crude lysates, we could reliably detect the expression of UcdO and UcdC proteins, though most UcdO was detected in the insoluble fraction (**Fig. 2b**). Nevertheless, incubation of urolithins (uroM6, uroM7, uroC, or uroA) with *ucd*-expressing crude lysates confirmed that only 9-position hydroxyl groups, present on uroM6 and uroC, were dehydroxylated to uroM7 and uroA, respectively (**Fig. 2c,d**). We did not observe any 10-position dehydroxylation in uroM6 or uroM7-treated lysates, demonstrating that the *E. asparagiformis ucd* operon specifically dehydroxylates 9-hydroxy urolithins. These results suggest that a different, unidentified operon is responsible for 10-position dehydroxylation in some *Enterocloster* spp.

**Figure 2.**
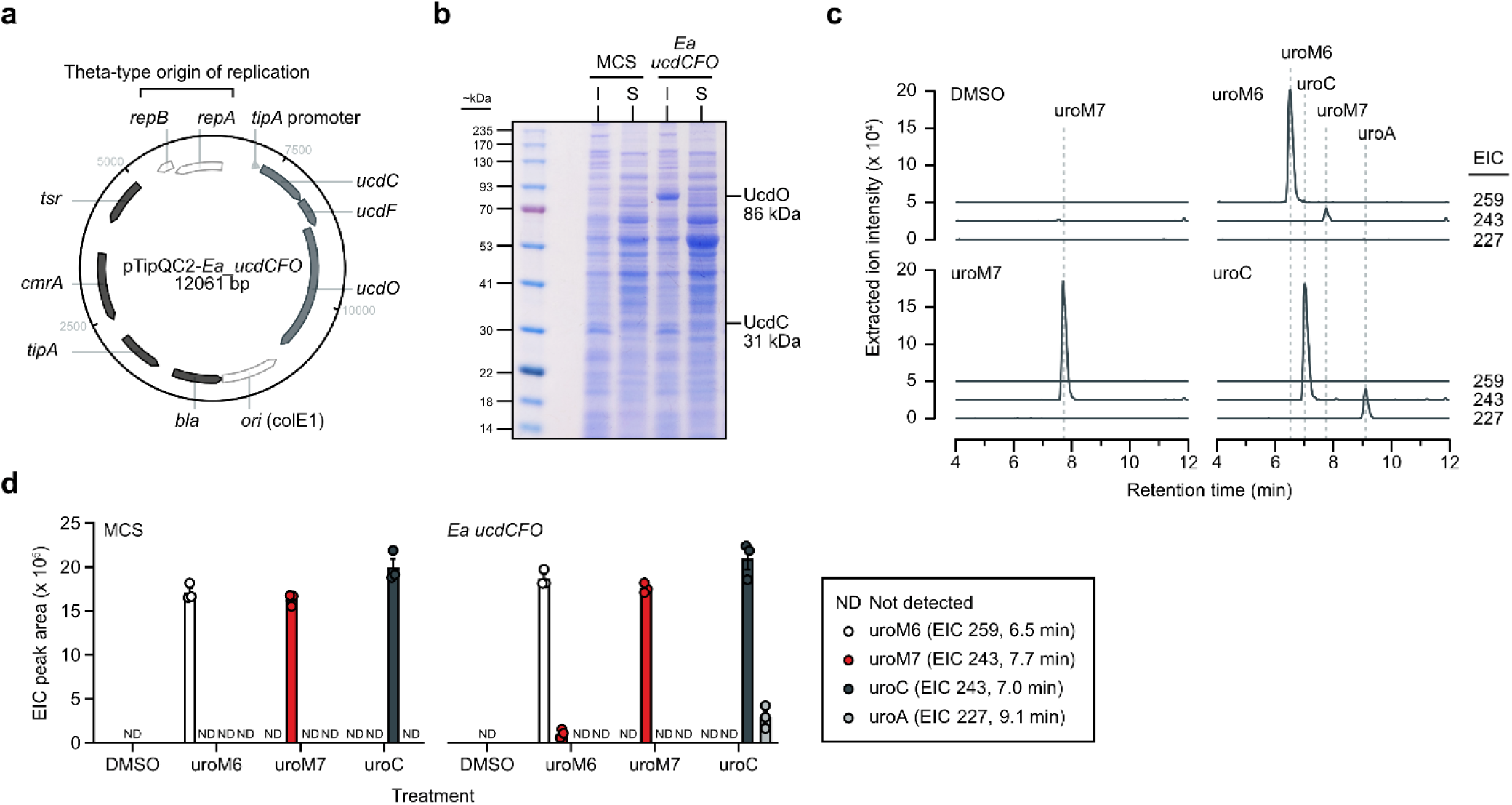
UroM6 is metabolized by distinct dehydroxylases in *Enterocloster* spp. **a** Map of the pTipQC2-*Ea_ucdCFO* plasmid for the heterologous expression of the *E. asparagiformis* (*Ea*) *ucd* operon in *R. erythropolis*. Plasmid map generated in Benchling. *ori*: origin of replication (*E. coli*); *bla*: beta-lactamase gene; *tipA*: thiostrepton-inducible transcriptional activator, *cmrA*: chloramphenicol efflux MFS transporter; *tsr*: thiostrepton resistance gene; *rep*: theta-type replicase (*R. erythropolis*). **b** Representative SDS-PAGE gel (10% bis-tris) of insoluble (I) and soluble (S) fractions from induced *R. erythropolis* pTipQC2-MCS and pTipQC2-*Ea_ucdCFO* crude lysates stained with colloidal Coomassie dye. **c** Aligned chromatograms (extracted ion intensities for [M-H]^-^: 259, 243, and 227) of DMSO or urolithin-treated crude lysates of *R. erythropolis* transformed with pTipQC2-*Ea_ucdCFO*. NADH (2.2 mM) was added to each lysate. Crude lysates were treated with urolithins (200 μM) for 48 h at room temperature anaerobically. Retention times of urolithin standards are shown with grey dotted lines. Data from n = 1 of 3 biological replicates are shown (related to (d)). **d** Quantification of urolithin extracted ion chromatogram (EIC) peak areas in crude lysates of *R. erythropolis* transformed with pTipQC2-MCS (multiple cloning site) or pTipQC2-*Ea_ucdCFO* plasmids (n = 3 biological replicates). Data are represented as means ± SEM.

### Ucd and Uxd dehydroxylases are induced by uroM6

Previous work from our lab has shown that dehydroxylation of urolithins by *Enterocloster* spp. is an inducible process triggered by substrate urolithins ^23^. Thus, we hypothesized that, upon exposure to uroM6 (**Fig. 1a**), *Enterocloster* spp. would express Ucd and <u>u</u>rolithin 10-position (<u>X</u>) <u>d</u>ehydroxylase (Uxd) proteins. Therefore, we individually incubated *E. asparagiformis*, *E. citroniae*, and *E. pacaense* with uroM6 (or DMSO as a vehicle control), pooled isolates by treatment group, and performed reference-based proteomics on pooled lysates (**Fig. 3a**). To ensure that target enzymes were being expressed, we collected samples that were actively metabolizing uroM6. In all 3 tested isolates, uroA could be detected after a 4 h incubation with uroM6 (**Fig. 3b**). Filtering proteins by presence/absence revealed that 44 individual proteins were overrepresented in the uroM6 group (**Fig. 3c**). Among these, the most abundant proteins induced by uroM6 mapped to *ucd* operon-encoded proteins (UcdC and UcdO) (**Fig. 3d**). However, we detected distinct xanthine dehydrogenase family proteins and FAD binding domain-containing proteins, which we annotated as putative Uxd proteins (**Fig. 3d**). The predicted FAD coenzyme-binding subunit (UxdC) was detected in all 3 species, while the substrate-binding oxidoreductase subunit (UxdO) was only detected in *E. pacaense*. Additionally, the molybdopterin cytosine dinucleotide (MCD) cofactor biosynthesis protein MogA (MOSC domain-containing protein) and the MCD cofactor chaperone protein XdhC (XdhC/CoxI family protein) ^26^ were expressed, though to lesser extents than Ucd and Uxd proteins (**Fig. 3d**).

**Figure 3.**
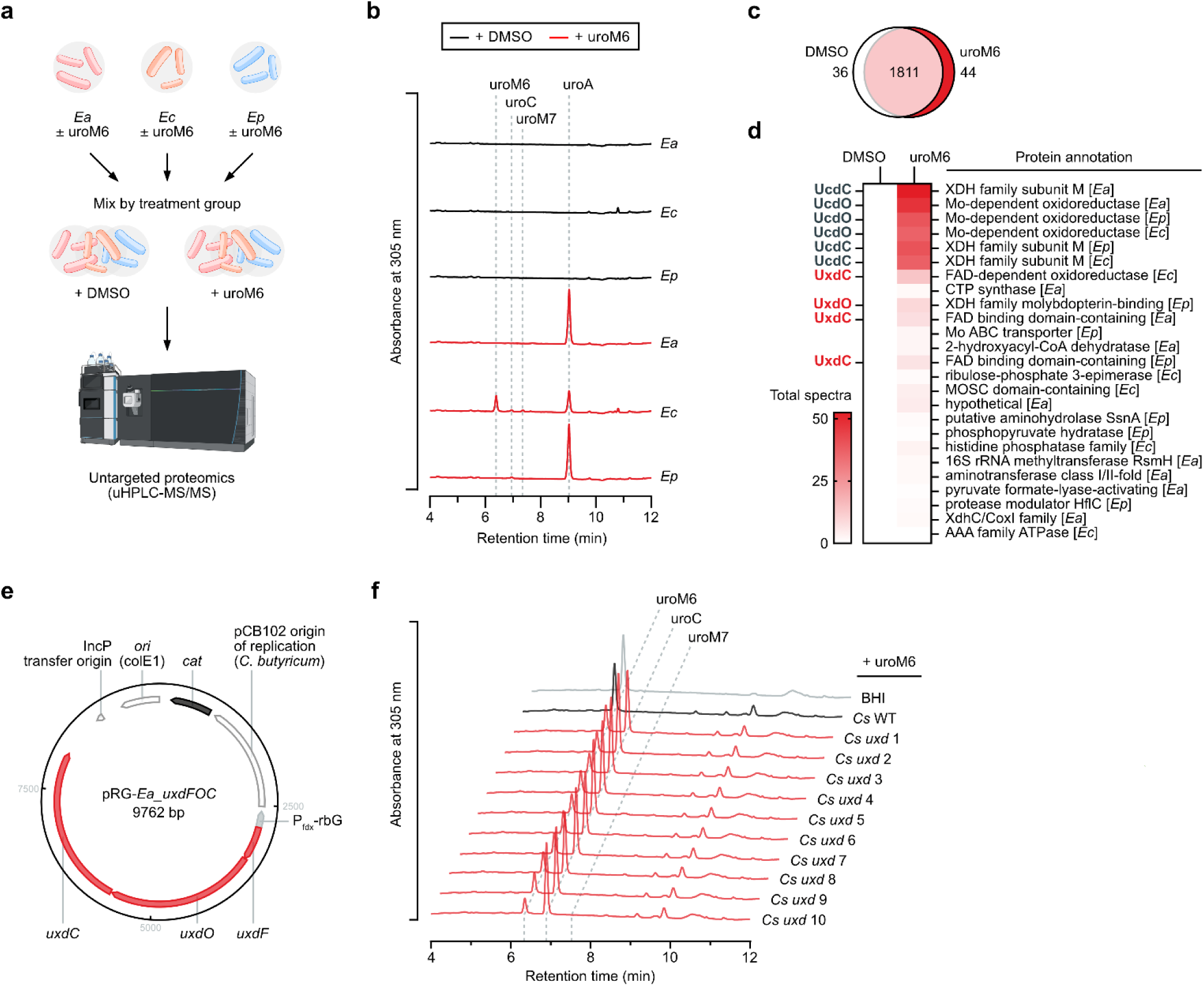
UroM6 induces the expression of two regioselective dehydroxylases. **a** Untargeted proteomics experimental design scheme. Treated isolates of *E. asparagiformis* (*Ea*), *E. citroniae* (*Ec*), and *E. pacaense* (*Ep*) were pooled by treatment group and lysed. Protein abundance was then determined by untargeted uHPLC-MS/MS. The scheme was generated using graphical elements from the NIAID NIH BIOART Source (bioart.niaid.nih.gov/bioart/42) and from BioRender. Castagner, B. (2025). **b** Aligned chromatograms (λ = 305 nm) of urolithins in DMSO or uroM6-treated (50 μM) mid-exponential phase isolates grown in mABB+H (4 h treatment with urolithins). Retention times of urolithin standards are shown with grey dotted lines. Data from n = 1 of 2 biological replicates are shown. **c** Venn diagram based on protein presence/absence in the untargeted proteomics dataset (from n = 2 biological replicates). **d** Heatmap of protein abundance (based on the average of total spectra from n = 2 biological replicates) for the 25 most abundant proteins detected in the uroM6 treatment group. Protein annotations from NCBI are shown on the right (along with the related taxon between brackets) and custom annotations are shown on the left. **e** Map of the pRG-*Ea_uxdFOC* plasmid for the heterologous expression of the *E. asparagiformis* (*Ea*) *uxd* operon in *C. sporogenes*. Plasmid map generated in Benchling. *ori*: origin of replication (*E. coli*); *cat*: chloramphenicol acetyltransferase; P_fdx_-rbG: promoter of the *C. sporogenes* ferredoxin gene with riboswitch G. **f** Offset chromatograms (λ = 305 nm) of urolithins in uroM6-treated (100 μM) mid-exponential phase *C. sporogenes* (*Cs*) cultures grown in BHI ± 30 μg/mL thiamphenicol (48 h treatment with uroM6). Retention times of urolithin standards are shown with grey dotted lines. Data from n = 1 biological replicate for each strain/transformant are shown. WT: wild type; *uxd* #: *Cs* pRG-*Ea_uxdFOC* clone number.

We next sought to validate the function of Uxd proteins through heterologous expression and noticed that, like Ucd, Uxd protein coding sequences were arranged in an operon with the following arrangement: *uxdFOC*. Therefore, we cloned the *E. asparagiformis uxd* operon into a shuttle plasmid to heterologously express UxdF, UxdO, and UxdC proteins in *R. erythropolis* (pTipQC2-*Ea_uxdFOC*). Unfortunately, electroporation of *R. erythropolis* with pTipQC2-*Ea_uxdFOC* did not yield any transformants capable of producing all three Uxd proteins, despite multiple attempts at transformation. We therefore searched the literature for alternative heterologous expression hosts that were both anaerobic and genetically tractable. Recent studies demonstrated that *Clostridium sporogenes*, found in anaerobic soil and gut environments, is an appropriate host for the heterologous expression of xanthine dehydrogenase family proteins ^27,28^. In addition, the RiboCas series of plasmids, which utilize theophylline-inducible riboswitches to control *Streptococcus pyogenes* Cas9 expression, have been used to delete the *spoIIE* gene in multiple *Clostridium* spp., including *C. sporogenes* ^29^. We therefore cloned the *E. asparagiformis uxd* operon and replaced the Cas9-encoding region of pRG-Cas2 to produce the pRG-*Ea_uxdFOC* shuttle plasmid (**Fig. 3e**). This plasmid was transformed into *C. sporogenes* via conjugation and resulting transformants were screened for protein expression and 10-position urolithin dehydroxylase activity. There were no obvious differences in protein expression between the wild type *C. sporogenes* and 10 randomly picked transformants (**Supplementary Fig. 2**). However, all 10 transformants were able to convert uroM6 to uroC, demonstrating 10-position urolithin dehydroxylase activity conferred by the *uxd* operon (**Fig. 3f**). Collectively, these data demonstrate that a subset of *Enterocloster* spp. express distinct enzymes (Ucd and Uxd), along with their accessory proteins, that regioselectively dehydroxylate uroM6.

### The *ucd* and *uxd* operons co-localize in urolithin-metabolizing *Enterocloster* spp. genomes

Since *ucd* and *uxd* operons appeared to be restricted to a subset of *Enterocloster* spp., we wondered about the prevalence of these operons both between and within *Enterocloster* spp. Therefore, we downloaded and queried 329 *Enterocloster* spp. genomes from the NCBI genomes database for the presence of *ucd* and *uxd* at the nucleotide level. As anticipated, the *ucd* operon was detected in *E. asparagiformis*, *E. bolteae*, *E. citroniae*, and *E. pacaense* genomes, whereas the *uxd* operon was detected in *E. asparagiformis*, *E. citroniae*, and *E. pacaense* (**Fig. 4a**). There was no evidence of these operons in other *Enterocloster* spp. genomes, including metagenome-assembled genomes (MAGs). All genomes of urolithin-metabolizing taxa encoded *uxd* and/or *ucd* with the exception of one *E. asparagiformis* MAG (GCA_040307595.1) which had neither of the operons (**Fig. 4a**). Altogether, *ucd* and *uxd* operons are highly prevalent within a subset of *Enterocloster* spp.

**Figure 4.**
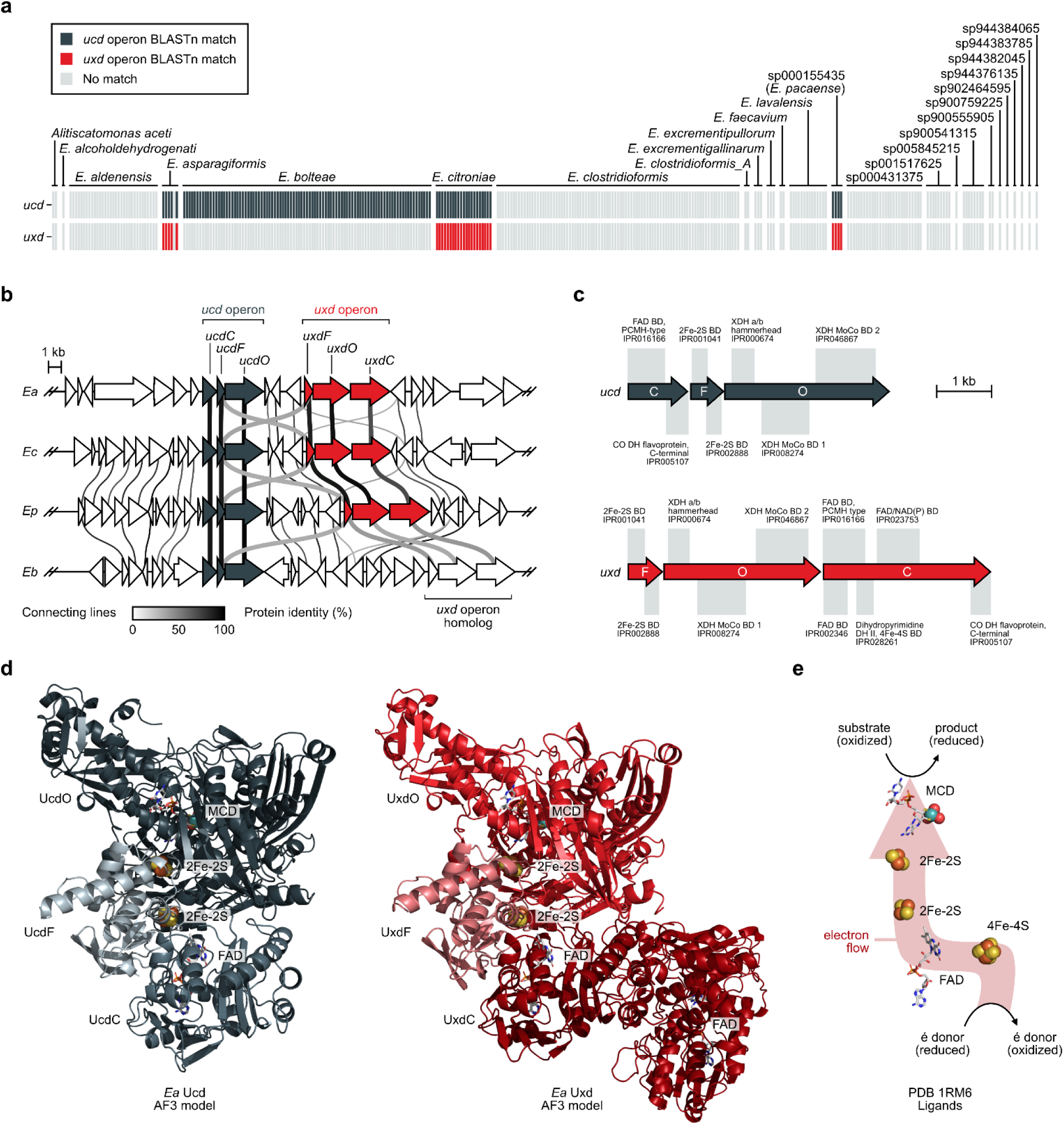
The *ucd* and *uxd* operons are in close proximity in *Enterocloster* spp. genomes. **a** Heat map of *ucd* (blue-grey) and *uxd* (red) operon presence based on BLASTn matches (% identity > 60 and length > 300) in *Enterocloster* spp. genomes downloaded from the NCBI. Genomes (n = 329) are grouped according to Genome Taxonomy Database (release 10-RS226) nomenclature. **b** Comparisons between *ucd* and *uxd* genomic contexts in urolithin-metabolizing *E. asparagiformis* (*Ea*), *E. citroniae* (*Ec*), *E. pacaense* (*Ep*), and *E. bolteae* (*Eb*). Alignments were performed using the clinker tool [https://doi.org/10.1093/bioinformatics/btab007] using genome accessions in **Supplementary Table 7**. Protein sequences with identities ≥ 60% are connected by lines. Bold lines indicate xanthine dehydrogenase family proteins within and between *ucd* and *uxd*-encoded proteins. **c** Representative domain annotations for *E. asparagiformis ucd* and *uxd*-encoded proteins. InterPro accessions are denoted below each annotation. FAD: flavin adenine dinucleotide; BD: binding domain; XDH: xanthine dehydrogenase; MoCo: molybdenum cofactor; CO: carbon monoxide; DH: dehydrogenase. **d** AlphaFold3 (AF3)-predicted protein structures of the *Ea* Ucd (shades of blue-grey) and Uxd (shades of red) protein complexes. A single trimer of the predicted heterohexamer is shown for clarity. FAD ligands were predicted by AF3. The molybdopterin cytosine dinucleotide cofactor (MCD) and 2Fe-2S clusters were derived from the superposition of AF3 structures onto the *Thauera aromatica* 4-hydroxybenzoyl-CoA reductase (PDB 1RM6). The 4Fe-4S cluster near the FAD (shown in **e**) was omitted as it is not common to both Ucd and Uxd. **e** Electron transport chain ligands in the *Thauera aromatica* 4-hydroxybenxoyl-CoA dehydrogenase enzyme complex (PDB 1RM6).

Next, we compared proteins in the vicinity of the *ucd* and *uxd* operons in urolithin-metabolizing *Enterocloster* spp. to investigate their genomic contexts. The *ucd* and *uxd* operons were in close proximity (<5 kb between *ucdO* and *uxdF*) in 10-position metabolizing species (*E. asparagiformis, E. citroniae,* and *E. pacaense*) (**Fig. 4b**). In these bacteria, Uxd proteins (UxdF, UxdO, and UxdC) were highly similar (>94, >96, and >83 % protein similarity, respectively) (**Supplementary Fig. 3a,b**). To our surprise, *E. bolteae*, which only metabolizes the 9-position of urolithins, possessed a *uxd* homolog near its *ucd* operon (**Fig. 4b**). In *E. bolteae*, the *uxd* operon homolog is more distant from the *ucd* (10.2 kb between *ucdO* and *uxdF*) and its Uxd proteins were less similar (<66 % protein similarity) compared to those from 10-position metabolizing species (**Supplementary Fig. 3a,b**). Notably, the substrate-binding UxdO protein was >96 % similar between *E. asparagiformis, E. citroniae*, and *E. pacaense*, but less than 63 % similar when compared to *E. bolteae*. These different protein sequences could explain the lack of 10-position dehydroxylase activity observed in *E. bolteae*.

We next compared domain annotations between *ucd* and *uxd*-encoded proteins of *E. asparagiformis*, which serves as our reference in this study. Iron-sulfur cluster-binding proteins (UcdF and UxdF) were similar in length, and both possessed two 2Fe-2S cluster binding domains, characteristic of prokaryotic xanthine dehydrogenase family proteins, which use these proteins to shuttle electrons between FAD and the MCD cofactor (**Fig. 4c**) ^30^. The MCD- and substrate-binding oxidoreductase subunits (UcdO and UxdO) both had functional domains typical of xanthine dehydrogenase family proteins: an XDH a/b hammerhead and molybdenum cofactor binding domains (**Fig. 4c**). The FAD-binding subunits (UcdC and UxdC) were more dissimilar, mostly due to the additional length of UxdC, which carried a 4Fe-4S cluster binding domain and an FAD/NAD(P) binding domain in addition to the FAD binding domain common to UcdC and UxdC (**Fig. 4c**). AlphaFold3 modeling ^31^ of Ucd and Uxd complexes produced similar overall structures resembling other xanthine dehydrogenase family proteins (**Fig. 4d**) ^32^. By superposing these predicted structures onto the crystal structure of the *Thauera aromatica* 4-hydroxybenzoyl-CoA reductase complex (dehydroxylase), we could place the MCD and 2Fe-2S clusters into UcdO and UxdO, and UcdF and UxdF, respectively. These ligands form an electron transport chain, likely enabling the dehydroxylation of substrates (e.g. urolithins) to their reduced products (**Fig. 4e**). In sum, *ucd* and *uxd* operons, which co-localize in urolithin-metabolizing *Enterocloster* spp. genomes, encode xanthine dehydrogenase proteins with similar predicted folds.

### Uxd expression is regulated by 9-hydroxy urolithins

We next wondered whether *uxd* operon substrates (10-hydroxy urolithins) induce the expression of *uxd* (**Fig. 5a**). Therefore, *E. asparagiformis*, grown to the mid-exponential phase, was treated with DMSO, uroM6, uroM7, uroC, or uroA (structures in **Fig. 1a**) and total RNA was extracted for RT-qPCR analysis focusing on *ucdO* and *uxdO* gene expression. Curiously, *uxd* expression followed the same pattern as *ucd* (**Fig. 5b**); that is, only 9-hydroxy urolithins (uroM6 and uroC) induced *uxd* expression. UroM7, which bears a hydroxyl group at the 10-position, and uroA, which is dehydroxylated at both the 9- and 10-positions, did not induce the expression of either operon (**Fig. 5b**). When we quantified urolithin concentrations in matched cultures (total RNA and urolithins isolated from the same samples), we observed dehydroxylation of uroM6 at both the 9- and 10-positions and dehydroxylation of uroC at the 9-position; however, uroM7 was unchanged (**Fig. 5c**). To validate that 9-hydroxy urolithins are required for 10-position dehydroxylation, *E. asparagiformis* cultures were induced with 9-hydroxy urolithins lacking a 10-hydroxyl group: urolithin D (uroD), uroC, or isouroA (**Fig. 5d**). After a 2 h incubation, 9-position dehydroxylation was observed for uroD- and uroC-treated cultures, yielding uroG and uroA, respectively; however, no dehydroxylation was observed for isouroA (at this timepoint) since uroB was not detected in culture supernatants (**Fig. 5e**). Following this 2 h induction, *E. asparagiformis* cells were washed and resuspended in PBS to minimize protein synthesis, then re-treated with urolithins to assess whether the 10-hydroxy urolithins could be dehydroxylated. In DMSO-induced resting cell suspensions, partial 9- and 10-position dehydroxylation was observed for uroM6 and uroC re-treated samples, suggesting basal activity or inducibility in these cell suspensions. Nevertheless, uroM7 remained unchanged (**Fig. 5f**), as previously observed in exponential phase cultures (**Fig. 5c**). In 9-hydroxy urolithin-induced resting cell suspensions, significant uroA production was observed for nearly all substrates (compared to DMSO-induced cultures). As anticipated, 10-position dehydroxylation of uroM7 to uroA was only observed in cell suspensions induced with 9-hydroxy urolithins (**Fig. 5f**), demonstrating the co-regulation of *ucd* and *uxd* operons in *E. asparagiformis*.

**Figure 5.**
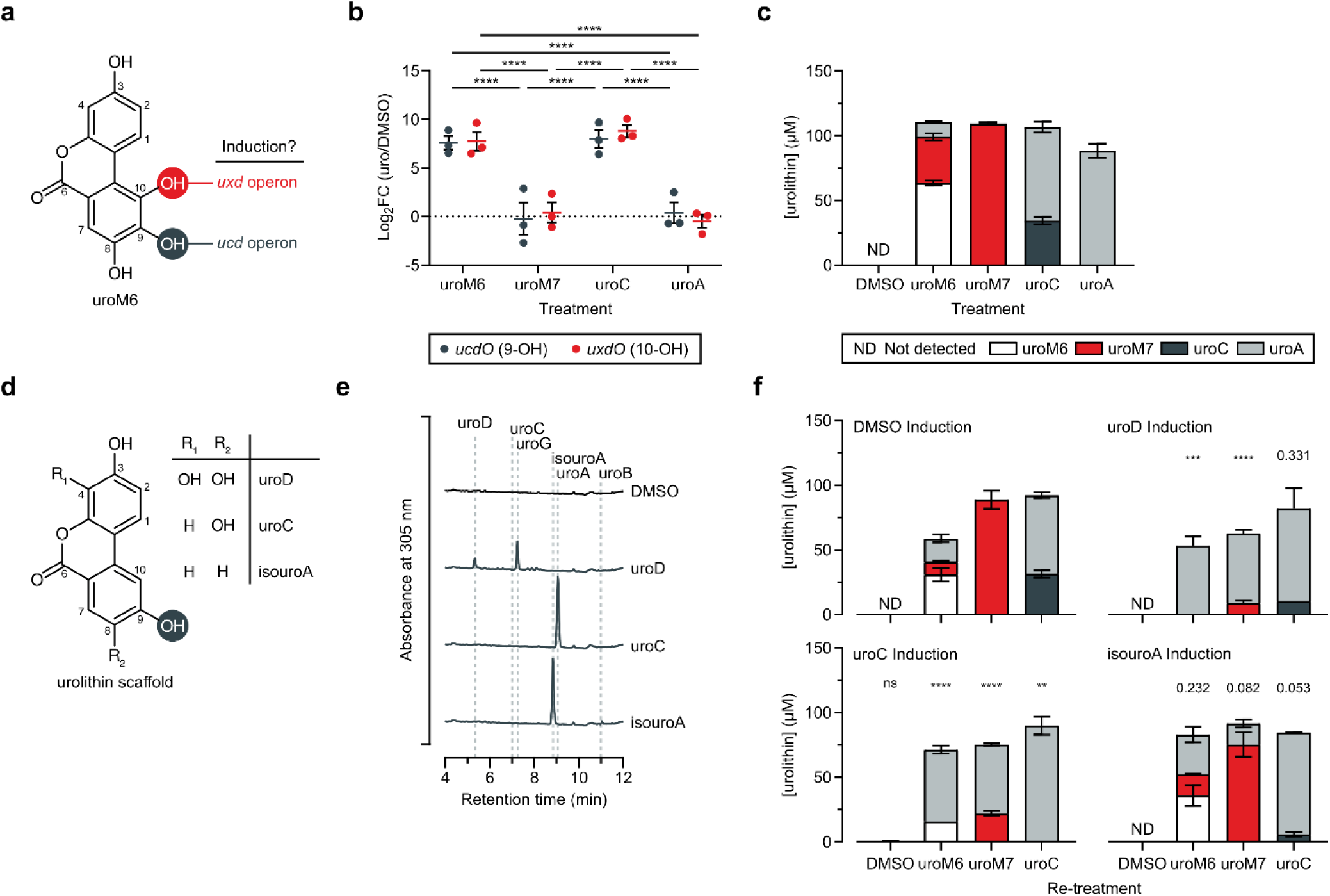
Transcription of *uxd* and *ucd* is coordinated. **a** Structure of urolithin M6 (uroM6). 9- and 10-position hydroxyl groups are highlighted in grey and red, respectively. **b** RT-qPCR expression of *E. asparagiformis* (*Ea*) *ucdO* and *uxdO* target after treatment with DMSO or urolithins (100 μM for 2 h in mABB+H media) (n = 3 biological replicates). Gene expression is displayed as log_2_FC (urolithin/DMSO); repeated-measures two-way ANOVA (matched by replicate and target gene) with Tukey’s multiple comparisons test. **c** Quantification of urolithin concentrations in matched samples from (**b**) determined by LC-MS (n = 3 biological replicates). **d** General urolithin structure with R groups showing hydroxyl group placement for urolithin D (uroD), urolithin C (uroC), and isourolithin A (isouroA). **e** Aligned chromatograms (λ = 305 nm) of DMSO or 9-hydroxy urolithin-induced (50 μM for 2 h) *E. asparagiformis* culture supernatants. Retention times of urolithin standards are shown with grey dotted lines. Data from n = 1 of 3 biological replicates are shown. **f** Quantification of urolithin concentrations in 9-hydroxy urolithin-induced (50 μM) *E. asparagiformis* cell suspensions re-treated with DMSO, uroM6, uroM7, or uroC (100 μM) (n = 3 biological replicates); two-way ANOVA (based on uroA concentration) with Dunnett’s multiple comparisons test (compared to re-treatment matched DMSO-induced samples). Bar colors are the same as in (**c**). Data in (**b**,**c**,**f**) are represented as means ± SEM. ** *P* < 0.01, *** *P* < 0.001, **** *P* < 0.0001.

In our previous study on the *ucd* operon, we performed RNA-seq experiments on DMSO and uroC-treated *E. asparagiformis* and *E. bolteae* ^23^. In *E. asparagiformis*, *uxd* operon genes were substantially induced by uroC (log_2_FC(uroC/DMSO) *Ea_uxdF*: 4.9, *Ea_uxdO*: 5.8, *Ea_uxdC*: 6.0) (**Supplementary Fig. 4a**); however, induction of the *E. bolteae uxd* homolog was less drastic (log_2_FC(uroC/DMSO) *Eb_uxdF*: 1.9, *Eb_uxdO*: 1.5, *Eb_uxdC*: 1.4) (**Supplementary Fig. 4b**). These differences in operon induction could also contribute to the lack of 10-position dehydroxylase activity observed in *E. bolteae*.

## Discussion

The human gut microbiota encodes millions of proteins with functions that remain largely unknown ^2,33^. This is notably the case for polyphenol-metabolizing enzymes (also termed polyphenol utilization proteins ^34^), which despite their widespread roles across ecosystems ^35,36^, remain poorly characterized and misannotated in reference databases ^37^. As urolithins can impact host health ^38^, understanding how these metabolites are produced can inform therapeutic microbiota modulation strategies ^38,39^. In this study, we used untargeted proteomics to uncover a 10-position urolithin dehydroxylase (Uxd) in a subset of urolithin-metabolizing *Enterocloster* spp. (*E. asparagiformis*, *E. citroniae*, and *E. pacaense*). Using diverse urolithins, which varied both in the number of hydroxyl groups and their positions on the parent structure, we characterized the regulation and substrate requirements for the induction and activity of urolithin dehydroxylases in *E. asparagiformis*.

The metabolism of dietary ellagitannins by the gut microbiota requires a diverse community of microorganisms harboring specialized enzymes (hydrolases, decarboxylases, and regioselective dehydroxylases) ^40^. We showed that EA-treated *Gordonibacter* spp. release urolithin intermediates in culture media, either through diffusion, leakage, or cell lysis ^41^, that become substrates for urolithin-metabolizing *Enterocloster* spp. In simple 2-member communities, *G. urolithinfaciens* paired with *E. bolteae* or *E. asparagiformis* could cooperate to metabolize EA to the terminal metabolite uroA (and to the intermediates uroM7 and uroG, respectively). These experiments also highlighted that *G. urolithinfaciens* cannot metabolize exogenous uroM6 (supplied directly), despite its ability to convert EA or uroM5 to uroC via uroM6 (**Fig. 1a**) ^42^. This observation could be explained by a lack of uroM6 transport or by a failure to induce Eadh2 (10-position dehydroxylase) in the ellagic acid metabolism (*eam*) gene cluster ^43^. Additionally, different urolithin intermediates accumulated in co-cultures depending on both their hydroxylation status upon release by *G. urolithinfaciens* and the dehydroxylases present in each *Enterocloster* spp. Indeed, along with uroA, substantial accumulation of uroM7 was observed with *E. bolteae*, whereas only some minor amounts of uroG were observed with *E. asparagiformis*. The abovementioned results indicate that urolithins could undergo substrate shunting whereby intermediates get trapped in alternative pathways, unable to be further dehydroxylated due to issues with transport or dehydroxylase induction. Additional studies are required to determine how the exchange of urolithin intermediates (cross-feeding) affects the fitness of urolithin-metabolizing species ^44^ and whether any species or strain level differences could alter the metabolic fate of urolithins in microbial communities.

Despite belonging to different phyla, both *Gordonibacter* spp. (Actinomycetota phylum) and a subset of urolithin-metabolizing *Enterocloster* spp. (Bacillota phylum) have evolved distinct molybdoenzymes to dehydroxylate the 10-position of urolithins (convergent evolution). The Eadh2 enzyme, part of the recently-described *eam* gene cluster in *G. urolithinfaciens*, belongs to the DMSO reductase family ^43^, while the Uxd complex belongs to the xanthine dehydrogenase family of enzymes. Most characterized polyphenol dehydroxylases belong to the DMSO reductase family ^45^. Examples of these enzymes include the *Eggerthella spp.* Hcdh (hydrocaffeic acid), Dadh (dopamine and L-norepinephrine), Cadh ((+)-catechin) and eCadh ((-)-epicatechin), and *Gordonibacter* spp. Hcdh (hydrocaffeic acid) and Dodh (3,4-dihydroxyphenylacetic acid (DOPAC)) ^45–47^. Interestingly, DMSO reductase family dehydroxylases require a *para*-hydroxyl group for dehydroxylation ^45,46^, whereas xanthine dehydrogenase family dehydroxylases, like Ucd, Uxd, and 4-HBCR, do not require this structural feature for enzyme activity ^23,48^. Indeed, isouroA (9-position to uroB), uroM7 (10-position to uroA), and 4-hydroxybenzoyl-CoA (4-position to benzoyl-CoA) are dehydroxylated at positions without a *para*-hydroxyl group by the abovementioned enzymes, respectively. These differences in molybdoenzyme family and substrate features suggest a different mechanism of uroM6 dehydroxylation for Eadh2 and Uxd ^43^.

Our comparative genomics analyses highlighted that a subset of urolithin-metabolizing *Enterocloster* spp. carry a *uxd* operon located in close proximity to their *ucd* operon. Surprisingly, *E. bolteae* possessed a *uxd* operon homolog in its genome. The endogenous substrate and function of this *uxd* homolog is currently unknown, though it is unlikely to dehydroxylate urolithins since *E. bolteae* only dehydroxylates urolithins at the 9-position via *ucd*. The evolutionary origins of the *ucd* and *uxd* operons are rather puzzling as both operons are co-regulated and encode xanthine dehydrogenase family proteins that dehydroxylate urolithins regioselectively, yet their genetic arrangement (*ucdCFO* vs. *uxdFOC*) and coenzyme-binding subunits differ substantially. Nevertheless, both AlphaFold3-predicted enzyme complexes adopted similar folds and subunit arrangements. Intriguingly, the UxdC coenzyme-binding subunit was predicted to bind 2 FAD molecules, at opposite ends, in agreement with the protein domain annotations. It is likely that UxdC utilizes 4Fe-4S clusters to shuttle electrons between the distal FAD towards the proximal FAD (next to UxdF), analogous to the role of the 4Fe-4S cluster in the *T. aromatica* 4-HBCR ^32^.

In this study, we focused on *Gordonibacter* and *Enterocloster* spp. since these gut-associated genera are both prevalent in human populations ^49^ and have been shown to cooperatively metabolize EA *in vivo* ^18^. It is worth noting that a growing list of bacterial strains have been reported to either partially or fully metabolize EA to terminal urolithin metabolites like uroA, isouroA, and uroB. These include *Ellagibacter isourolithinifaciens* ^20^, *Streptococcus thermophilus* FUA329 ^50^, *Lactococcus garvieae* FUA009 ^51^, *Enterococcus faecium* FUA027 ^52^, *Limosilactobacillus fermentum* FUA033 ^53^, and *Bifidobacterium pseudocatenulatum* INIA P815 ^54^. Unfortunately, with the exception of *E. isourolithinifaciens* ^43^, many of these strains lack publicly accessible genomes and are generally not commercially available, making polyphenol-metabolizing enzyme discovery and validation difficult. Future enzyme discovery efforts using these strains, and others yet to be discovered, would enable a better understanding of the diversity of enzymes involved in EA and urolithin metabolism across taxa in the gut microbiota.

The transcription of gut bacterial xenobiotic metabolizing enzymes is often an inducible process triggered upon exposure to their substrates ^55,56^. In urolithin-metabolizing *Enterocloster* spp., the *ucd* operon (9-position dehydroxylase) is induced upon treatment with 9-hydroxy urolithins like uroM6, uroC, and isouroA ^23^. Surprisingly, *uxd* transcription was not triggered by uroM7 (3,8,10-trihydroxy-urolithin) and no 10-position dehydroxylation was observed when this compound was supplied directly. Instead, *ucd* induction by its substrates was required for *uxd* transcription and for the conversion of uroM7 to uroA. These findings suggest that the *ucd* and *uxd* operons, which are separated by less than 3 kb in *E. asparagiformis* genome, share regulatory proteins that coordinate their transcription, along with accessory proteins for MCD cofactor biosynthesis, after sensing 9-hydroxy urolithins. This regulatory mechanism is consistent with the urolithin metabolites that are released by EA-metabolizing *Gordonibacter* spp. (uroM5, uroM6, and uroC), which all feature a 9-position hydroxyl group. Thus, *Gordonibacter* spp. and *Enterocloster* spp. may be particularly well adapted to cooperatively metabolize EA in the gut, as reflected by their complementary urolithin-metabolizing enzymes.

In sum, our study highlights the genes, proteins, and substrate features underlying urolithin metabolism in a subset of *Enterocloster* spp. Dietary polyphenols and their postbiotic metabolites are gaining widespread attention for their roles in host health ^57^, though many of molecular mechanisms underlying their production by gut bacteria remain unknown. Enzyme discovery efforts described here and elsewhere ^45,47,58,59^ not only contribute to our basic understanding of polyphenol-bacteria ecology but also explain some of the inter-individual differences in polyphenol metabolism (metabotypes ^7^) observed in human populations.

## Materials and Methods

### General reagents

A complete list of reagents and chemicals used in this study is provided in **Supplementary Table 1**. Reference numbers for commercial media, kits, and master mixes are provided in relevant sections below.

### Anaerobic bacterial strains and culturing conditions

Strains of bacteria used in this study are listed in **Supplementary Table 2**. *Enterocloster* spp. and *Gordonibacter* spp. were grown from glycerol stocks on mABB+H (recipe below) or BHI (Difco #237500) agar plates for 48-72 h at 37 °C in a Coy vinyl anaerobic chamber, which was maintained with a gas mixture of 3% H_2_, 10% CO_2_, 87% N_2_. To make overnight cultures, a single isolated colony was inoculated into 5 mL of mABB+H or BHI liquid medium for *Enterocloster* spp. and BHI of BHIrf (BHI + 1% (w/v) arginine + 10mM sodium formate) ^47^ for *Gordonibacter* spp. Cultures were incubated at 37 °C between 16-24 h for *Enterocloster* spp. and 48 h for *Gordonibacter* spp.

### Modified anaerobe basal broth with hemin (mABB+H)

For 1 L of modified anaerobe basal broth (mABB, based on Oxoid Anaerobe Basal Broth), the following components were dissolved in MilliQ water, then autoclaved: 16 g peptone, 7 g yeast extract, 5 g sodium chloride, 1 g starch, 1 g D-glucose monohydrate, 1 g sodium pyruvate, 0.5 g sodium succinate, 1 g sodium thioglycolate, 15 g agar (for plates). The autoclaved solution was allowed to cool, then the following filter-sterilized solutions were added aseptically: 10 mL of 100 mg/mL L-arginine-HCl, 10 mL of 50 mg/mL L-cysteine, 8 mL of 50 mg/mL sodium bicarbonate, 50 μL of 10 mg/mL vitamin K1, 20 mL of 50 mg/mL dithiothreitol, and, for mABB+H, 10 mL of 0.5 mg/mL haemin. The media was then placed in the anaerobic chamber and allowed to reduce for at least 24 h prior to its use in experiments.

### Genomic DNA extraction of isolates and identity validation

Genomic DNA was extracted from 0.5-1 mL of overnight culture using the One-4-All Genomic DNA Miniprep Kit (BioBasic #BS88503) according to the manufacturer’s instructions. The identities of all bacteria in this study were validated by full-length (V1-V9) 16S rRNA sequencing using primers 16S_V1_27_f and 16S_V9_1492_r listed in **Supplementary Table 3**. The purified genomic DNA was used as a template for PCR reactions using the Q5 High-Fidelity polymerase (NEB #M0491). PCR tubes were placed in a thermal cycler and targets (∼1.5 kb) were amplified according to the following cycling conditions: 30 s at 98 °C, 30 cycles (10 s at 98 °C, 20 s at 60 °C, 45 s at 72 °C), 2 min at 72 °C, and hold at 10 °C. 5 μL of the reaction was mixed with 6X Gel Loading Dye (NEB #B7024A) and loaded onto a 1% agarose gel (made with 1X TAE buffer) containing FroggaStain (FroggaBio #FB555). PCR product sizes were compared to the Quick-Load Purple 1 kb Plus DNA Ladder (NEB #N0550).

PCR products (∼1.5 kb) were purified using the Monarch PCR & DNA Cleanup Kit (NEB #T1030) according to the manufacturer’s instructions for products < 2 kb. Purified 16S PCR products were eluted in nuclease-free water, quantified using the Qubit dsDNA HS assay kit (Invitrogen #Q32851), and adjusted to 30 ng/μL. Linear/PCR product sequencing was performed by Plasmidsaurus using Oxford Nanopore Technology with custom analysis and annotation.

### *Gordonibacter* spp. ellagic acid metabolism assays

Overnight cultures of *Gordonibacter* spp. (**Supplementary Table 2**) were diluted 1/50 in fresh BHIrf media. Ellagic acid (10 mM stock in DMSO) or an equal volume of DMSO were added to a final concentration of 100 μM. Cultures were incubated at 37°C under anaerobic conditions and samples were collected after 2 days. For urolithin detection in supernatants, 300 μL of culture was centrifuged at 5000 x g for 3 min and the supernatant was collected. Metabolites in cultures and supernatants were extracted following *Extraction Method C*.

### *Enterocloster* spp. treatments with urolithins

Urolithins used in this study (**Supplementary Table 1**) were dissolved in DMSO to a concentration of 10 mM. Overnight cultures of *Enterocloster* spp. (**Supplementary Table 2**) were diluted 1/50 into fresh media and incubated at 37 °C in an anaerobic chamber. After 5-7 hours (5 h for *E. asparagiformis* and *E. pacaense,* and 7 h for *E. citroniae*) of incubation, 10 mM urolithins (or an equivalent volume of DMSO) were added to the growing cultures at a final concentration of 50 or 100 μM for protein expression or RNA expression, respectively. For protein expression analyses, the cultures were incubated for an additional 4 h. For RNA expression analyses and inducibility tests, the cultures were incubated for an additional 2 h. Metabolites in cultures were extracted following *Extraction Method A*.

### Co-cultures of *G. urolithinfaciens* and *Enterocloster* spp

Overnight cultures of *G. urolithinfaciens* (*Gu*), *E. bolteae* (*Eb*), and *E. asparagiformis* (*Ea*) were diluted 1/100 (each) into fresh BHI media (5 mL) according to the following (4) treatment groups: BHI (sterile), *Gu*, *Gu* + *Eb*, *Gu* + *Ea*. Each treatment group was split into 3 × 500 µL aliquots and treated with DMSO, ellagic acid (10 mM stock in DMSO), or urolithin M6 (10 mM stock in DMSO) to a final concentration of 100 μM. Samples were incubated at 37 °C under anaerobic conditions and 200 μL samples were taken after 5 days and stored at −70°C until metabolite extraction. Metabolites in cultures were extracted following *Extraction Method A*.

### Urolithin extraction from bacterial cultures

Frozen (−70 °C) bacterial cultures were thawed at room temperature. For quantification of urolithin concentrations, urolithin standards (stock 10 mM in DMSO) were spiked into separate media aliquots immediately before extraction.

#### Extraction Method A

This method was used for cultures and their supernatants. Samples were diluted with an equal volume of MeOH, vortexed briefly, and incubated at 50 °C for 10 min. Samples were centrifuged at 20,000 x g for 5 min to pellet insoluble material, then transferred to LC-MS vials. Urolithins were then analyzed by LC-MS.

#### Extraction Method B

This method was used to extract urolithins from crude bacterial lysates. Lysates were diluted with 3 volumes of MeOH, vortexed briefly, and incubated at room temperature for 10 min. Samples were centrifuged at 20,000 x g for 5 min to pellet insoluble material, then transferred to LC-MS vials. Urolithins were then analyzed by LC-MS.

#### Extraction Method C

This method was used for cultures and supernatants. 3 volumes of ethyl acetate with 0.1% formic acid were added to the sample and vortexed briefly. Samples were centrifuged at 10,000 x g for 5 min to separate the organic and inorganic layers. The organic layer was collected and dried using a rotary evaporator. Concentrated samples were dissolved in 50% methanol, centrifuged at 20,000 x g for 5 min to pellet insoluble material, then transferred to LC-MS vials. Urolithins were then analyzed by LC-MS.

### LC-MS method to quantify urolithins

Samples were injected (10 μL) into a 1260 Infinity II Single Quadrupole LC/MS system (Agilent) fitted with a Poroshell 120 EC-C18 4.6×50 mm, 2.7 μm column (Agilent #699975-902) with a compatible guard column (Agilent #820750-911) incubated at 30 °C. The mobile phase was composed of MilliQ water + 0.1% formic acid (solvent A) and acetonitrile + 0.1% formic acid (solvent B). The flow rate was set to 0.7 mL/min. The gradient was as follows: 0-8 min: 10-30 %B, 8-10 min: 30-100 %B, 10-13.5 min: 100 %B isocratic, 13.5-13.6 min: 100-10 %B, then 13.6-15.5 min: 10 %B. The multiple wavelength detector was set to monitor absorbance at 305 nm. The mass spectrometer (API-ES) was run in negative mode in both selected ion monitoring (SIM) and scan (100-1000 m/z) modes to validate peak identities. Capillary voltage was set to 3000 V, drying gas to 10.0 L/min, nebulizer pressure to 30 psig, and drying gas temperature to 350 °C. Peaks were validated based on retention times compared to spike-in standards and mass-to-charge ratios. To quantify urolithins, integrated peak areas (derived from Agilent ChemStation software) for the compounds of interest were compared to spike-in standards of known concentrations. When standards were not available or overlapped with other peaks, the extracted ion chromatogram peak area was used: ellagic acid ([M-H]^-^: 301), urolithin M5 ([M-H]^-^: 275), urolithin E ([M-H]^-^: 259), urolithin D ([M-H]^-^: 259), urolithin M7 ([M-H]^-^: 243), urolithin M6 ([M-H]^-^: 259), urolithin C ([M-H]^-^: 243), urolithin G ([M-H]^-^: 243), urolithin A ([M-H]^-^: 227), isourolithin A ([M-H]^-^: 227), urolithin B ([M-H]^-^: 211). Blank runs of 50% MeOH were included at the beginning and end of LC-MS sequences to ensure proper column washing.

### Plasmid construction in *E. coli*

All plasmids used in this study were constructed through DNA assembly using the NEBuilder HiFi DNA Assembly Master Mix (NEB #E2621). Linearized pTipQC2 plasmid DNA (Hokkaido Systems Science Co. #RE-0006) was generated through overnight (∼16 h) restriction digestion with NdeI (NEB #R0111) and XhoI (NEB #R0146) in rCutSmart buffer at 37 °C. Linearized pRG_BB (backbone) plasmid DNA was generated through PCR amplification of the pRG-Cas2 plasmid (RiboCas plasmid series ^29^) using the Q5 DNA polymerase (NEB # M0491). To reduce contamination from the pRG-Cas2 template, DpnI (NEB #R0176) was added to the pRG_BB PCR reaction after thermal cycling and incubated for 1 h at 37 °C. DNA inserts were generated through PCR amplification of *E. asparagiformis* genomic DNA using the Q5 DNA polymerase. In all cases, Q5 DNA polymerase PCR cycling was as follows: 30 s at 98 °C, 30 cycles (10 s at 98 °C, 20 s at T_a_, reaction-specific extension time at 72 °C), 2 min at 72 °C. Primer sequences are listed in **Supplementary Table 3** and specific PCR conditions (primers, T_a_, and extension time) are listed in **Supplementary Table 4**. Restriction digest products and PCR amplicons were loaded into 0.6-1% (w/v) agarose gels (in 1X TAE) and purified using the Monarch DNA Gel Extraction Kit (NEB #T1020) according to the manufacturer’s instructions for different DNA sizes. Purified DNA concentrations were determined using the Qubit dsDNA HS assay kit (Invitrogen #Q32851) and fragments were mixed for HiFi DNA assembly at 50 °C for 1 h according to the manufacturer’s instructions. HiFi DNA assembly constructs (plasmid backbones and inserts) are provided in **Supplementary Table 5**.

HiFi DNA assembly reactions were diluted ¼ in nuclease-free water and 2 μL were transformed into homemade CaCl_2_ competent *E. coli* NEB10β (NEB #C3019) via heat shock. Cells were spread onto selective LB + 100 μg/mL ampicillin agar plates and incubated at 37 °C overnight. The next day, colonies were picked and grown in selective LB + 100 μg/mL ampicillin. Plasmids were purified using the Plasmid DNA Miniprep Kit (BioBasic #BS414) and size was confirmed with a diagnostic restriction digest. Whole plasmid sequencing was performed by Plasmidsaurus using Oxford Nanopore Technology with custom analysis and annotation. GenBank accessions for sequenced plasmids are listed in **Supplementary Table 3**. Plasmid maps were generated in Benchling (2025): https://benchling.com.

### pTipQC2 plasmid transformation into *R. erythropolis*

Electrocompetent *R. erythropolis* were prepared by inoculating 50 mL NBYE (0.8% w/v nutrient broth, 0.5% yeast extract) + 0.05% Tween-80 media with 1 mL of a stationary phase (48-72 h growth from a single colony) *R. erythropolis* culture and grown aerobically for 16 h at 30 °C with shaking at 200 RPM. The next day, cells were pelleted at 5,000 x g for 10 min at 4 °C and washed according to the following sequence: 2 washes of (10 mL of ice cold sterile MilliQ water), 10 mL of ice cold sterile 10% glycerol. The final pellet was resuspended in 2.5 mL of ice cold sterile 10% glycerol. The resuspended electrocompetent *R. erythropolis* were aliquoted (50 μL/aliquot), flash-frozen in liquid nitrogen, and stored at −80 °C long-term. For transformations, frozen electrocompetent cells were diluted in sterile MilliQ water, then 3 μL (∼0.5-1 μg) of various plasmid DNA (isolated from *E. coli*) were added to appropriate tubes. Cells with plasmid DNA were transferred to 0.1 cm gap cuvettes (Bio-Rad #165-2089) and electroporated (1.8 kV, 25 μF, 200 Ω). Time constants were between 4.0-4.4 ms. The cuvette was immediately filled with 1 mL LB post shock and cells were allowed to recover at 30 °C for 3 h before plating 100 μL of undiluted culture and of the remaining concentrated recovery culture on LB + 30 μg/mL chloramphenicol plates at 30 °C. After 3-4 days of incubation, colonies were picked and grown in selective liquid LB + 30 μg/mL chloramphenicol at 30 °C with shaking at 200 RPM. Ucd protein-producing colonies were verified by inducing protein expression and assaying proteins by SDS-PAGE (described below).

### Heterologous expression of Ucd proteins in *R. erythropolis*

All growth steps below were performed in selective media (LB + 30 μg/mL chloramphenicol) in aerobic conditions at 30 °C with shaking at 220 RPM. Single colonies of *R. erythropolis* transformed with pTipQC2-MCS (multiple cloning site) or pTipQC2-*Ea_ucdCFO* were inoculated into 15 mL selective media and grown for 72 h to produce overnight cultures. Overnight cultures were then thoroughly resuspended and diluted 1:10 into 25 mL fresh selective media and grown for 7-8 h until OD_600_ values reached ∼0.6. Thiostrepton (5 mg/mL in DMSO) was added to a final concentration of 0.5 μg/mL and cultures were incubated aerobically for 16 h at 30 °C to induce protein expression. The next morning, 20 mL from each culture were pelleted and resuspended in 4 mL of lysis buffer (1 X PBS, 1 mM DTT, 1% Triton X-100, 2 mg/mL egg white lysozyme, and 1 tablet/10 mL cOmplete Mini, EDTA-free (Roche #11836170001)). The resuspended cells in lysis buffer were incubated on ice for 1 h with shaking, then sonicated on ice using a Misonix Sonicator 3000 set to power level 3/10 according to the following sequence (aerobically, in a cold room): 20 s ON, 40 s OFF, for a total of 4 min ON. To evaluate dehydroxylase activity, crude lysates (100 μL) were treated with 2 μL of 10 mM urolithin solution (or DMSO vehicle) and with 8 μL of 30 mM NADH (dissolved in lysis buffer), then incubated anaerobically at chamber temperature for 48 h. Metabolites in lysates were extracted following *Extraction Method B*.

### SDS-PAGE analysis of Ucd proteins

Crude lysates described above were centrifuged for 2 min at 20,000 x g. The insoluble pellet was separated from the soluble supernatant. The insoluble pellet (from 100 μL of crude lysate) was resuspended in 100 μL of 1X reducing loading dye (62.5 mM Tris-HCl (pH 6.8), 2% (w/v) SDS, 10% glycerol, 0.01% (w/v) bromophenol blue, 38 mM DTT). The soluble fraction was diluted with 4X reducing loading dye to a final concentration of 1X. All samples were heated at 95 °C for 5 min, then 15 μL were loaded onto a 10% bis-tris polyacrylamide protein gel. Gels were fixed and stained with 10X colloidal Coomassie solution (0.4% (w/v) Coomassie Brilliant Blue R250, 10% (w/v) citric acid, 8% (w/v) ammonium sulfate, and 20% (v/v) methanol).

### Protein extraction from uroM6-treated *Enterocloster* spp

All steps other than sonication were carried out under anaerobic conditions. To extract proteins, 10 mL of treated (50 μM urolithin M6 for 4 h) *Enterocloster* spp. culture (see *Enterocloster spp. treatments with urolithins*) were pelleted (6,500 g for 3 min) and the supernatant was discarded. The pellet was washed with 10 mL of pre-reduced PBS, pelleted again, and resuspended in 0.6 mL of pre-reduced lysis buffer (20 mM Tris, pH 7.5, 500 mM NaCl, 10 mM MgSO_4_, 10 mM CaCl_2_, and 1 tablet/10 mL cOmplete Mini, EDTA-free (Roche #11836170001)). The resuspended pellet was then sonicated on ice using a Misonix Sonicator 3000 set to power level 2/10 according to the following sequence (aerobically, in a cold room): 20 s ON, 40 s OFF, for a total of 2 min ON. Tubes were centrifuged at 20,000 x g for 2 min to pellet insoluble particles and the supernatant was kept for downstream analyses.

### Proteomics analysis of uroC-treated *Enterocloster* spp

Extracted proteins were submitted for proteomic analysis at the RI-MUHC. For each sample, protein lysates were loaded onto a single stacking gel band to remove lipids, detergents, and salts. The gel band was reduced with DTT, alkylated with iodoacetic acid, and digested with trypsin. Extracted peptides were re-solubilized in 0.1% aqueous formic acid and loaded onto a Thermo Acclaim Pepmap (Thermo, 75 um ID X 2 cm C18 3 um beads) precolumn and then onto an Acclaim Pepmap Easyspray (Thermo, 75 um ID X 15 cm with 2 um C18 beads) analytical column separation using a Dionex Ultimate 3000 uHPLC at 250 nL/min with a gradient of 2-35% organic (0.1% formic acid in acetonitrile) over 3 hours. Peptides were analyzed using a Thermo Orbitrap Fusion mass spectrometer operating at 120,000 resolution (FWHM in MS1) with HCD sequencing (15,000 resolution) at top speed for all peptides with a charge of 2+ or greater. The raw data were converted into *.mgf format (Mascot generic format) for searching using the Mascot 2.6.2 search engine (Matrix Science) against a custom *Enterocloster* spp. protein database composed of *Enterocloster asparagiformis* (GenBank Assembly GCA_025149125.1), *Enterocloster citroniae* (GenBank Assembly GCA_001078435.1), *Enterocloster pacaense* (GenBank Assembly GCA_900566185.1), and a database of common contaminant proteins. Mascot was searched with a fragment ion mass tolerance of 0.100 Da and a parent ion tolerance of 5.0 ppm. O-63 of pyrrolysine, carboxymethyl of cysteine and j+66 of leucine/isoleucine indecision were specified in Mascot as fixed modifications. Deamidation of asparagine and glutamine and oxidation of methionine were specified in Mascot as variable modifications.

The database search results were loaded into Scaffold Q+ Scaffold_5.0.1 (Proteome Sciences) for statistical treatment and data visualization. Scaffold (v5.3.0) was used to validate MS/MS based peptide and protein identifications. Peptide identifications were accepted if they could be established at greater than 95.0% probability by the Peptide Prophet algorithm ^60^ with Scaffold delta-mass correction. Protein identifications were accepted if they could be established at greater than 99.0% probability and contained at least 2 identified peptides. Protein probabilities were assigned by the Protein Prophet algorithm ^61^. Proteins that contained similar peptides and could not be differentiated based on MS/MS analysis alone were grouped to satisfy the principles of parsimony. Proteins sharing significant peptide evidence were grouped into clusters. Protein quantification and differential expression were determined in Scaffold using the following parameters: Quantitative method was set to total spectra and the minimum value was set to 0.5 in case proteins were not detected in one condition.

### pRG plasmid conjugation into *C. sporogenes*

Homemade CaCl_2_ competent *E. coli* CA434 conjugal donor cells (carrying the R702 plasmid) ^62^ were transformed with the pRG-*Ea_uxdFOC* plasmid via heat shock, allowed to recover in SOC medium (BioBasic #SD7009), and plated onto LB + 30 μg/mL chloramphenicol (for the pRG plasmids) + 15 μg/mL tetracycline (for the R702 plasmid). Plates were incubated at 30 °C aerobically.

The day before conjugation, single colonies of CA434 donor cells carrying pRG-*Ea_uxdFOC* were inoculated into 5 mL of LB + 30 μg/mL chloramphenicol + 15 μg/mL tetracycline and grown overnight at 30 °C with shaking (aerobically). A single colony of the recipient *C. sporogenes* was inoculated into 5 mL of BHI and grown overnight at 37 °C (anaerobically). The next day, 1.5 mL of CA434 donor was pelleted at 6,500 x g for 3 min, washed with 1 mL of sterile PBS, re-pelleted, placed on ice, and brought to the anaerobic chamber. An aliquot from the *C. sporogenes* overnight culture (in BHI) was taken out of the anaerobic chamber and heat shocked at 50 °C for 5 min to increase transformation efficiency ^63^. Afterwards, 300 μL of heat shocked *C. sporogenes* recipient culture was mixed with the pellets from appropriate donors. The resulting mixture was spotted (3 x 20 μL) onto pre-reduced BHI plates (without antibiotics) and incubated for 24 h at 37 °C (anaerobically). The following day, all 3 spots were scraped with a 10 μL sterile loop and resuspended in 300 μL of PBS. 100 μL of this suspension was spread onto a BHI agar plate supplemented with 250 μg/mL D-cycloserine (against CA434) + 30 μg/mL thiamphenicol (for pRG-*Ea_uxdFOC*) and grown for 24 h at 37 °C. Individual transformants were picked and re-streaked onto BHI + 250 μg/mL D-cycloserine + 30 μg/mL thiamphenicol. Plasmid-positive *C. sporogenes* colonies were identified by colony PCR (cPCR) using the OneTaq 2X Quick-Load Master Mix (NEB #M0486) with *cat*_cPCR primers and *Ea_uxdC*_cPCR_f/RepOri_cPCR_r and conditions detailed in **Supplementary Tables 3,4**. In all cases, PCR cycling was as follows: 5 min at 94 °C, 35 cycles (20 s at 94 °C, 30 s at T_a_, reaction-specific extension time at 68 °C), 5 min at 68 °C. Cultures from pRG-*Ea_uxdFOC*-positive colonies, grown in BHI + 30 μg/mL thiamphenicol, were mixed with 50% glycerol and stored at −70 °C.

### Heterologous expression of Uxd proteins and dehydroxylase activity assay in *C. sporogenes*

All growth steps below were performed in selective media (BHI + 30 μg/mL thiamphenicol) in anaerobic conditions at 37 °C. Single colonies of *C. sporogenes* transformed with pRG-*Ea_uxdFOC*, grown on BHI + 30 μg/mL thiamphenicol agar plates, were inoculated into 5 mL selective media and grown overnight. Overnight cultures were then diluted 1/100 into 10 mL fresh selective media and grown for 6 h. Theophylline (100 mM in 0.1 M NaOH) was added to a final concentration of 2 mM to induce protein expression. For the initial screen, the culture was split into 200 μL aliquots that were then treated with 100 μM uroM6. Whole cell cultures were incubated for 48 h at 37 °C (anaerobically). To evaluate dehydroxylase activity, urolithin metabolites were extracted from whole cultures following *Extraction Method A*.

### SDS-PAGE analysis of Uxd proteins

*C. sporogenes* pRG-*Ea_uxdFOC* cultures induced with 2 mM theophylline for 24 h were pelleted at 5,000 g for 5 min and the pellets were resuspended in 1.5 mL of PBS + 2 mg/mL lysozyme. The cell suspensions were sonicated on ice using a Misonix Sonicator 3000 set to power level 2/10 according to the following sequence (aerobically, in a cold room): 20 s ON, 40 s OFF, for a total of 3 min ON. The crude lysate was diluted with 4X reducing loading dye to a final concentration of 1X (62.5 mM Tris-HCl (pH 6.8), 2% (w/v) SDS, 10% glycerol, 0.01% (w/v) bromophenol blue, 38 mM DTT). All samples were heated at 95 °C for 5 min, then 10 μL were loaded onto a 10% bis-tris polyacrylamide protein gel. Gels were fixed and stained with 10X colloidal Coomassie solution (0.4% (w/v) Coomassie Brilliant Blue R250, 10% (w/v) citric acid, 8% (w/v) ammonium sulfate, and 20% (v/v) methanol).

### Comparative genomics

The analysis of *ucd* and *uxd* operon presence in *Enterocloster* spp. genomes was done by first downloading all available (n = 452) *Enterocloster* spp. genome sequences (FASTA format) from the NCBI genomes database (excluding atypical genomes) [https://www.ncbi.nlm.nih.gov/datasets/genome/?taxon=2719313&typical_only=true] (Accessed 01-08-2025). All FASTA files with GenBank GCA_#########.# accessions wereretained for further analysis (n = 452). The Genome Taxonomy Database (GTDB) table for *Enterocloster* spp. (release 10-RS226 (16-04-2025)) was downloaded from [https://gtdb.ecogenomic.org/searches?s=al&q=Enterocloster] (Accessed 02-08-2025). Each genome was queried using rBLAST (v1.2.0, DOI: 10.18129/B9.bioc.rBLAST) for matches to the *E. bolteae ucd* operon and *E. asparagiformis uxd* operon nucleotide sequences with criteria: - perc_identity 60 -max_target_seqs 5 -subject_besthit. Matches were further filtered for length > 300 bases to exclude small fragments. NCBI GenBank accessions for all queries without GTDB taxonomy or the "Undefined (Failed Quality Check)" flag were removed from further analyses, yielding a total of n = 329 genomes (accessions listed in **Supplementary Table 6**). Presence/absence of each operon was plotted in ggplot2 (v.3.5.2). All code, tables, and resulting RData files from this analysis were deposited in Zenodo [https://doi.org/10.5281/zenodo.16740616].

Comparisons between *ucd* and *uxd* operon genomic contexts were done using CAGECAT ^64^ (v1.0, https://cagecat.bioinformatics.nl/) using the clinker tool for visualization. GenBank files of *ucd* and *uxd* operon genomic contexts were downloaded from the NCBI Sequence Viewer from genome accessions in **Supplementary Table 7**. The identity threshold was set to 60% and proteins were manually recolored to highlight conserved features. Protein similarity matrices were generated from clinker alignment.txt output files.

### Protein domain annotations

Domain annotations for *E. asparagiformis* Ucd proteins (UcdC, UcdF, and UcdO) and Uxd (UxdF, UxdO, UxdC) proteins were obtained from InterPro ^65^ via UniProt accessions listed in **Supplementary Table 8**.

### Protein structures and homology modeling

Protein structures were generated using AlphaFold3 (AF3) [https://alphafoldserver.com/] with protein sequences from *E. asparagiformis* (*Ea*) listed in **Supplementary Table 8**. For the *Ea* Ucd complex, a single copy of UcdC, UcdF, UcdO protein sequences and a single FAD ligand were used as input. For the *Ea* Uxd complex, a single copy of UxdC, UxdF, UxdO protein sequences and 2 FAD ligands were used as input. AF3 output model_0.cif (crystallographic information file) files were opened in PyMOL (v2.4.1, Schrodinger LLC). For homology modeling, FoldSeek ^66^, integrated into the AlphaFold Protein Structure Database [https://alphafold.ebi.ac.uk/], was used to search the RCSB Protein Data Bank (RCSB PDB) [https://www.rcsb.org/] for proteins that had similar folds to the *Ea* UcdO and UxdO subunits. The crystal structure of the *Thauera aromatica* 4-hydroxybenzoyl-CoA reductase (PDB 1RM6) ^32^ was used as a reference to position (using the “super” command in PyMOL to align oxidoreductase subunits) the molybdopterin cytosine dinucleotide (MCD) cofactor and 2Fe-2S clusters in UcdO/UxdO and UcdF/UxdF, respectively.

### RNA extraction and RT-qPCR

A volume of 1 mL of treated (DMSO or 100 μM uroM6, uroM7, uroC, or uroA for 2 h) *E. asparagiformis* culture was pelleted (6,500 g for 3 min) and the supernatant was removed for later LC-MS analysis. The pellet was then mixed with 900 μL TRI reagent (Zymo Research #R2050-1-50) and transferred to a ZR BashingBead lysis tube (Zymo Research #S6012-50). Samples were lysed in a Mini Beadbeater 16 (Biospec #607) according to the following sequence: 1 min ON, 5 min OFF. Bead beating was done for a total of 2 min ON to preserve longer transcripts. RNA isolation was then performed using the Direct-zol RNA Miniprep Kit (Zymo Research # R2051) according to the manufacturer’s instructions (including an on-column DNase digestion). To ensure complete DNA removal, an additional DNA digestion step was performed on the isolated RNA using the Ambion DNA-free DNA Removal Kit (Invitrogen #AM1906) according to the manufacturer’s instructions. RNA concentration and quality were verified by NanoDrop and 1% (w/v) agarose gel electrophoresis.

Isolated RNA samples (500 ng) were reverse transcribed using the iScript Reverse Transcription Supermix (Bio-Rad #1708840) in a reaction volume of 10 μL. The iScript No-RT Control Supermix was used as a no enzyme control for reverse transcription (-RT). The reaction mixtures were incubated in a thermal cycler: 5 min at 25 °C, 20 min at 48 °C, and 1 min at 95 °C. Both cDNA and -RT controls were diluted 1/20 in nuclease-free water before use. qPCR reactions were conducted using the Luna Universal qPCR Master Mix kit (NEB #M3003). The *Ea_ucdO*_qPCR, *Ea_uxdO*_qPCR, and *Ea*_CTP_Ref primer pairs (**Supplementary Table 3**) were added to their respective master mixes (final primer concentration of 250 nM) and 7 μL of diluted template (cDNA, -RT, no template) were added to 28 μL of master mix for technical triplicates. Replicate mixes were pipetted (10 μL/well) into a MicroAmp Fast 96-Well Reaction Plate (Applied Biosystems #4346907) and the plates were sealed (PlateSeal #PS-PETSTXL-100), then spun down for 2 min to eliminate air bubbles. The qPCR detection parameters on the Viia7 qPCR machine (Applied Biosystems #4346907) were as follows: SYBR Green detection, ROX reference dye, 10 μL reaction volume. The thermal cycling conditions were: 1 min at 95 °C, 40 cycles (15 s at 95 °C, 30 s at 60 °C), and melt analysis (60-95 °C). Data were analyzed according to the 2^-ΔΔCt^ method with the *E. asparagiformis* CTP synthase (*Ea*_CTP) gene serving as the reference gene due to its stable expression upon treatment with uroC ^23^.

### *E. asparagiformis* cell suspension assays to test dehydroxylase inducibility

Overnight cultures of *E. asparagiformis* (*Ea*) were diluted 1/50 into fresh mABB+H media (25 mL) and grown for 5 h at 37 °C in anaerobic conditions. Each culture was then split into 4 × 5 mL aliquots, and each aliquot was treated with DMSO or 9-hydroxy urolithins (uroD, uroC, or isouroA) (50 μM final concentration for each compound) for 2 at 37 °C to induce protein expression. Cultures were then pelleted at 6,500 x g for 3 min and supernatants were collected for downstream extraction and LC-MS analysis using *Extraction Method A*. The pellets were washed with 5 mL of pre-reduced PBS, pelleted once again, and resuspended in 1 mL of PBS. This cell suspension was split into 4 × 200 μL aliquots, and each aliquot was treated with either DMSO, uroM6, uroM7, or uroC (100 μM final concentration). Samples were briefly vortexed and incubated at room temperature in the anaerobic chamber for 20 h. Metabolites in cultures were extracted following *Extraction Method A*.

### Statistical analyses and graphing

Statistical methods were not used to determine sample sizes, experiments were not randomized, and the investigators were not blinded. Data points were assumed to be normally distributed, though this was not formally tested. Details related to each statistical test performed are supplied in the Figure legends and statistical test results are provided in the Source data file for each figure panel, where applicable. In all cases, α = 0.05 and tests were two-tailed. Data were plotted in GraphPad Prism (v10.4.0) or ggplot2 (v3.5.2). Figures were assembled in Affinity Designer 2 (v2.6.3.332).

## Data availability

Untargeted proteomics datasets generated in this study were deposited in the PRIDE repository under accession PXD066865 and DOI 10.6019/PXD066865 (Reviewer access username: and password: dtegALAUbFOQ). DNA sequences of plasmids generated in this study will be deposited in GenBank upon publication: pTipQC2-*Ea_ucdCFO* (#XYZ), pTipQC2-*Ea_uxdFOC* (#XYZ), and pRG-*Ea_uxdFOC* (#XYZ).

RNA-sequencing data for uroC-treated *E. bolteae* and *E. asparagiformis* are available at NCBI SRA BioProject ID PRJNA996126 under BioSample accession codes SAMN36514640 and SAMN36514641, respectively. The *T. aromatica* 4-HBCR X-ray crystal structure is available in the Protein Data Bank (PDB) under accession 1RM6.

## Code availability

All code, tables, and RData files obtained from the analysis of *Enterocloster* spp. genomes were deposited in Zenodo [https://doi.org/10.5281/zenodo.16740616].

## Acknowledgements

This research was funded by the Canadian Institutes of Health Research (CIHR) grant PJT-437944, a Weston Family Microbiome Initiative’s 2021 Transformational Research program, and a Weston Family Foundation 2024 Proof-of-Principle grant to B.C. B.C. held a tier II Canada Research Chair (CRC) in Therapeutic Chemistry (2015-2025). R.P. is supported by the CIHR Canada Graduate Scholarship-Doctoral (#493808 for 2023-2026) and by the Fonds de Recherche du Québec-Santé : Bourse de formation au doctorat (#316063 for 2022-2023). A.G. is supported by the CIHR Canada Graduate Scholarship-Masters for 2024-2025.

Untargeted proteomics sample preparation and analysis was performed by the Research Institute of the McGill University Health Centre (RI-MUHC) Proteomics and Molecular Analysis Platform. Molecular biology experiments were enabled by the McGill University Imaging and Molecular Biology Platform (IMBP).

## Author information

### Contributions

**R.P.** designed the study, performed experiments, analyzed data, created figures, and wrote the initial manuscript. **A.G.** performed experiments, analyzed data, and created figures. **L.D.** consulted on experimental design and methodology. **B.C.** designed the study, supervised the research, obtained research funding, and wrote the initial manuscript with R.P. All authors reviewed and edited the manuscript.

## Supplementary Figures

**Supplementary Figure 1.**
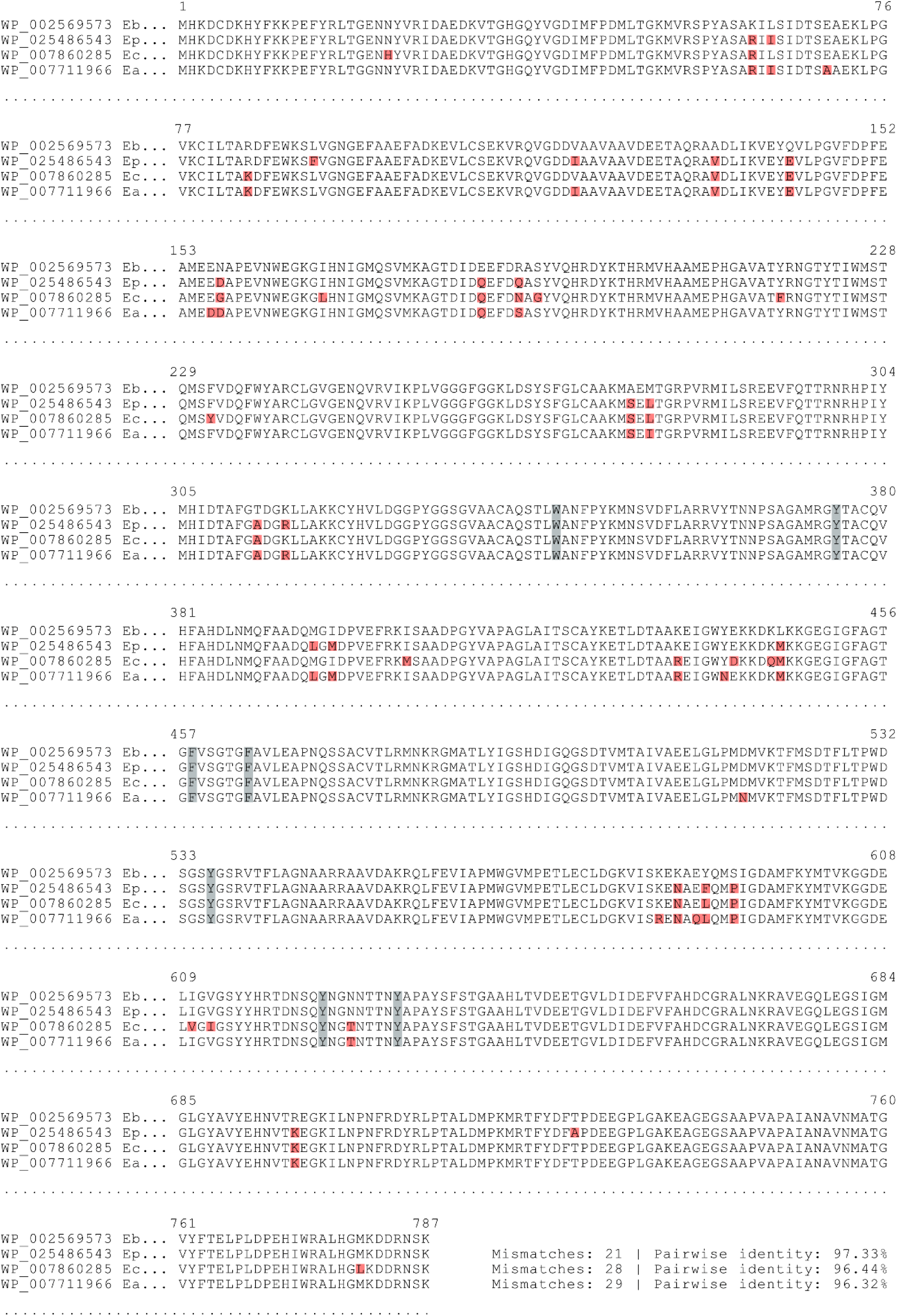
*Enterocloster* spp. UcdO protein multiple sequence alignment. Clustal Omega multiple sequence alignment of *Enterocloster* spp. UcdO sequences (787 AA) generated in Benchling. Residues boxed in red represent mismatches relative to the *E. bolteae* (*Eb*) UcdO template and residues boxed in grey represent predicted urolithin binding site residues (W345, Y375, F458, F464, Y536, Y624, and Y632). The number of mismatches and % pairwise identity (relative to *Eb*) are provided at the end of each sequence. NCBI GenBank accessions are provided on the left of each line and in **Supplementary Table 8**.

**Supplementary Figure 2.**
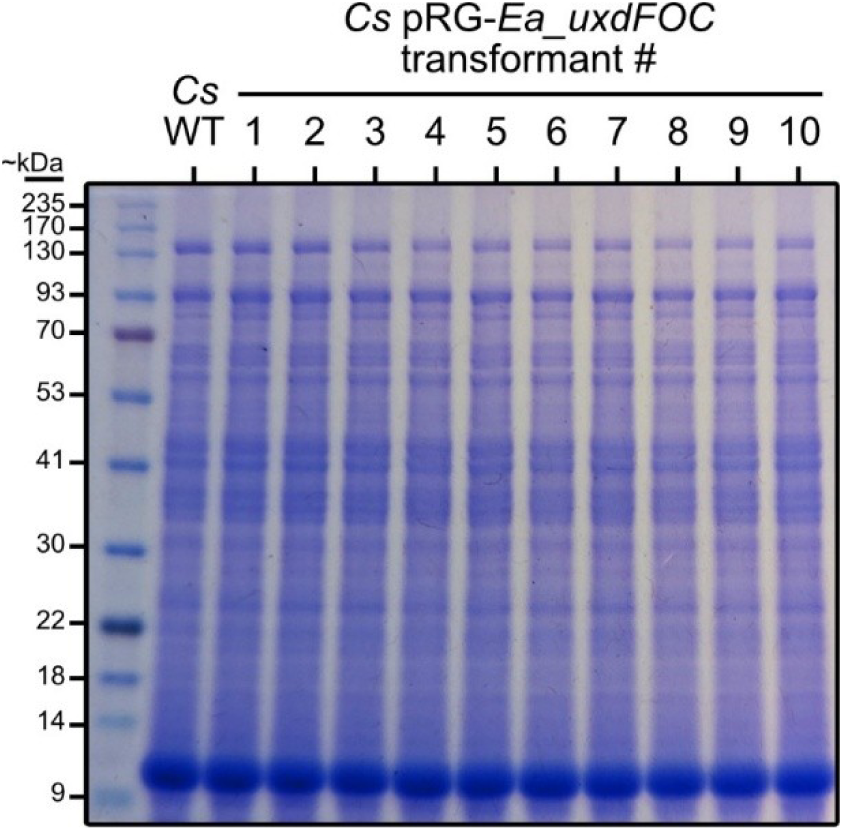
SDS-PAGE analysis of *C. sporogenes* pRG-Ea_uxdFOC lysates. Representative SDS-PAGE gel (10% bis-tris) of total lysates from 2 mM theophylline-induced *C. sporogenes* wild type (WT) and *C. sporogenes* pRG-*Ea_uxdFOC* transformants stained with colloidal Coomassie dye.

**Supplementary Figure 3.**
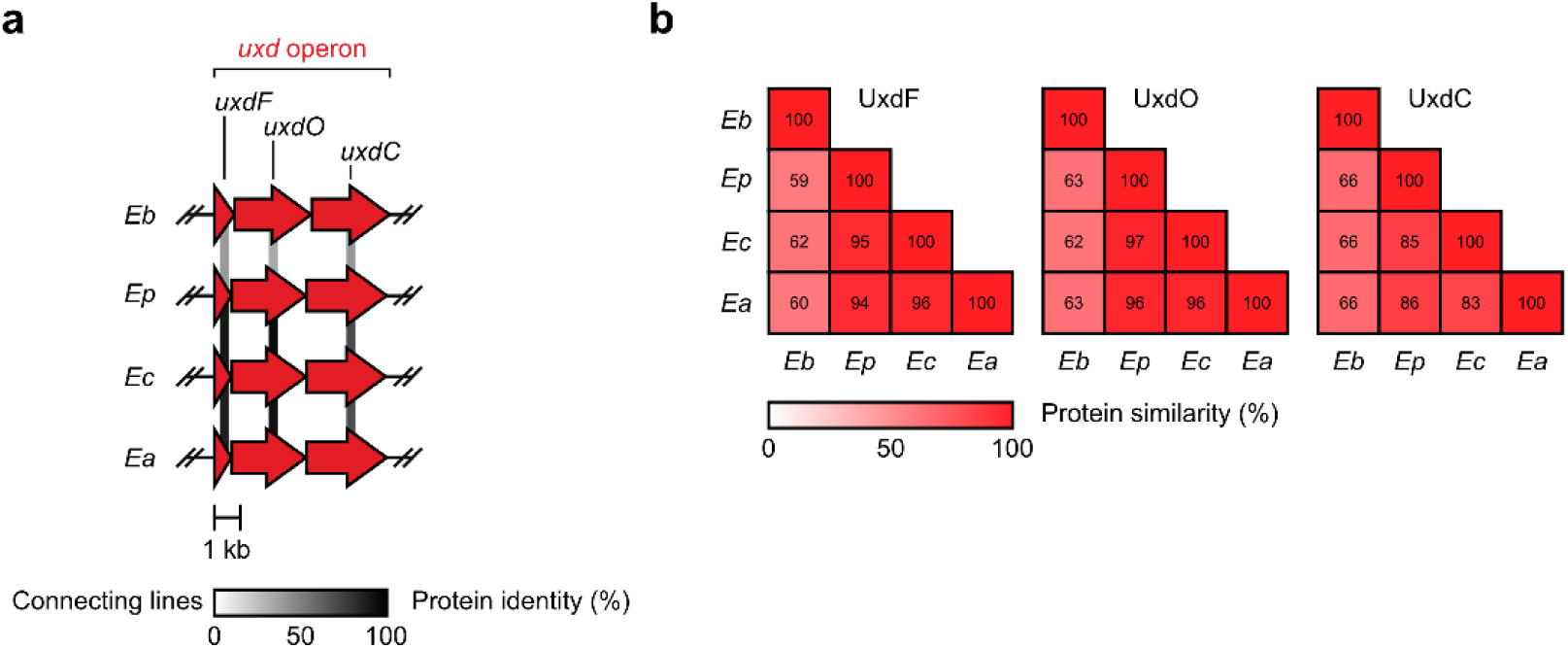
*Enterocloster* spp. Uxd protein multiple sequence alignment. **a** Multiple sequence alignment of *Enterocloster* spp. *uxd* operon-encoded proteins performed in clinker ^64^. Lines connecting *uxd* genes indicate pairwise protein identities. **b** Protein similarity matrix for each Uxd protein. NCBI GenBank assemblies used as input for clinker are provided in **Supplementary Table 7**.

**Supplementary Figure 4.**
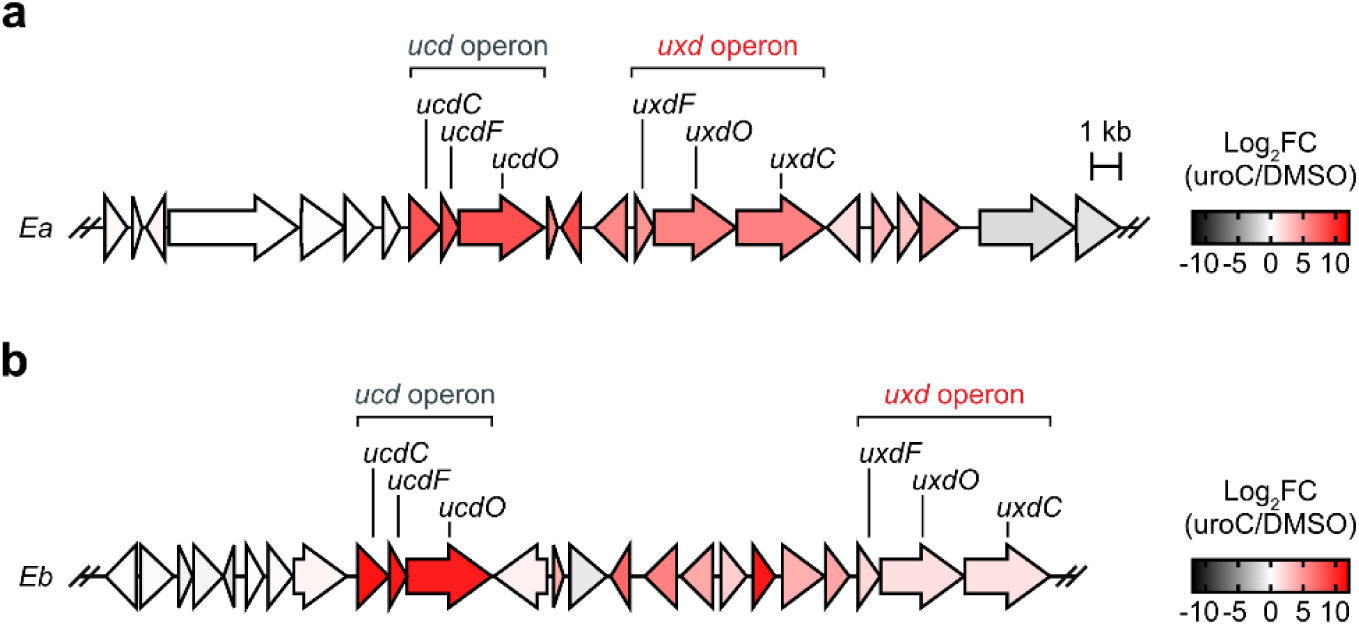
Urolithin C-treated *Enterocloster* spp. *ucd* and *uxd* differential expression (RNA-seq). **a** Differential gene expression (RNA-seq) of genes of interest in uroC (100 μM)-treated *E. asparagiformis* (*Ea*). Data from n = 4 biological replicates. **b** Differential gene expression (RNA-seq) of genes of interest in uroC (100 μM)-treated *E. bolteae* (*Eb*). Data from n = 4 biological replicates. Log_2_FC values were obtained from data in ^23^ derived from RNA-sequencing reads described in the Data Availability section.

## Supplementary Tables

**Supplementary Table 1.**
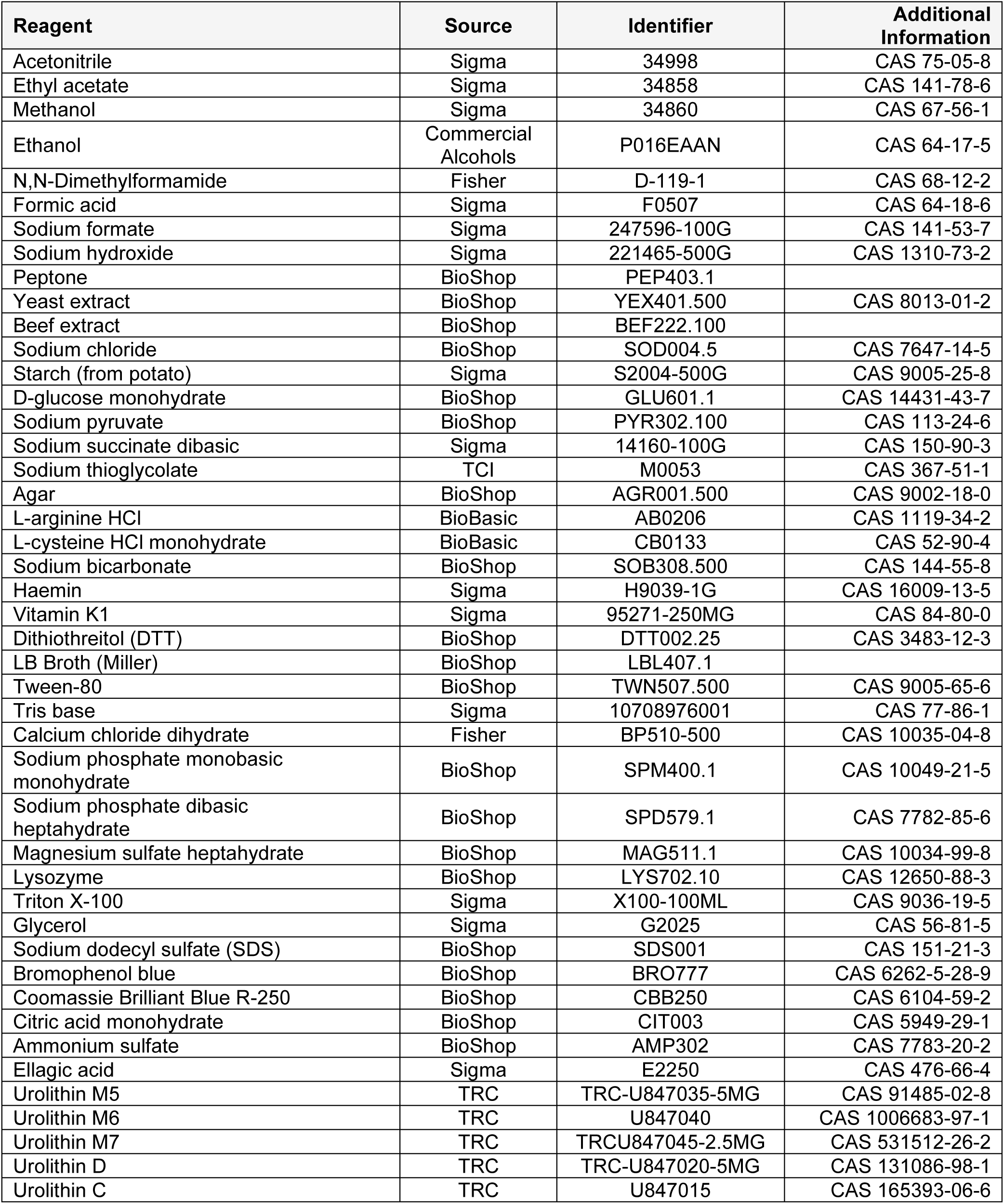
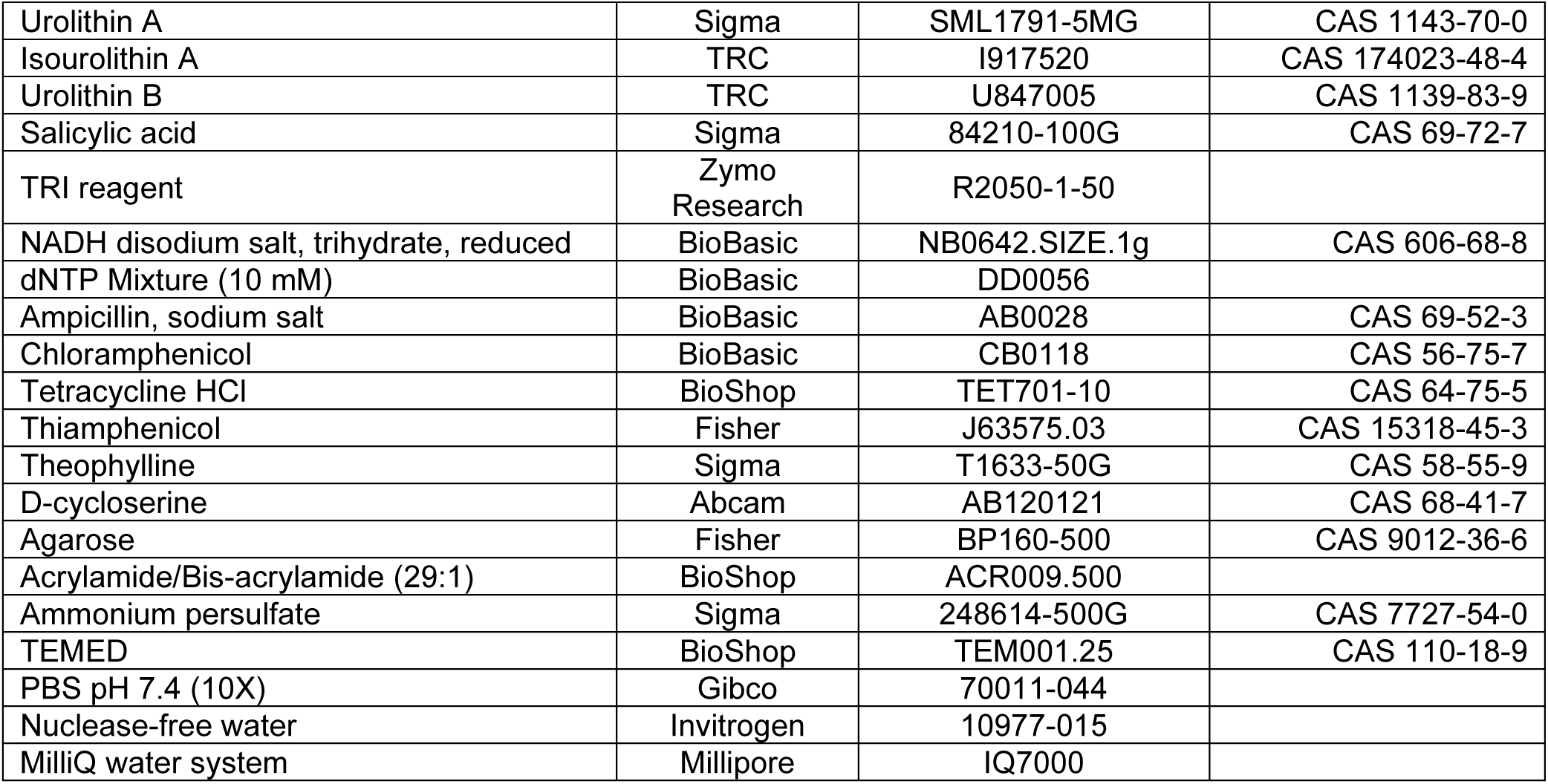
Reagent and chemical reference numbers.

| Reagent | Source | Identifier | Additional Information |
| --- | --- | --- | --- |
| Acetonitrile | Sigma | 34998 | CAS 75-05-8 |
| Ethyl acetate | Sigma | 34858 | CAS 141-78-6 |
| Methanol | Sigma | 34860 | CAS 67-56-1 |
| Ethanol | Commercial Alcohols | P016EAAN | CAS 64-17-5 |
| N,N-Dimethylformamide | Fisher | D-119-1 | CAS 68-12-2 |
| Formic acid | Sigma | F0507 | CAS 64-18-6 |
| Sodium formate | Sigma | 247596-100G | CAS 141-53-7 |
| Sodium hydroxide | Sigma | 221465-500G | CAS 1310-73-2 |
| Peptone | BioShop | PEP403.1 |  |
| Yeast extract | BioShop | YEX401.500 | CAS 8013-01-2 |
| Beef extract | BioShop | BEF222.100 |  |
| Sodium chloride | BioShop | SOD004.5 | CAS 7647-14-5 |
| Starch (from potato) | Sigma | S2004-500G | CAS 9005-25-8 |
| D-glucose monohydrate | BioShop | GLU601.1 | CAS 14431-43-7 |
| Sodium pyruvate | BioShop | PYR302.100 | CAS 113-24-6 |
| Sodium succinate dibasic | Sigma | 14160-100G | CAS 150-90-3 |
| Sodium thioglycolate | TCI | M0053 | CAS 367-51-1 |
| Agar | BioShop | AGR001.500 | CAS 9002-18-0 |
| L-arginine HCl | BioBasic | AB0206 | CAS 1119-34-2 |
| L-cysteine HCl monohydrate | BioBasic | CB0133 | CAS 52-90-4 |
| Sodium bicarbonate | BioShop | SOB308.500 | CAS 144-55-8 |
| Haemin | Sigma | H9039-1G | CAS 16009-13-5 |
| Vitamin K1 | Sigma | 95271-250MG | CAS 84-80-0 |
| Dithiothreitol (DTT) | BioShop | DTT002.25 | CAS 3483-12-3 |
| LB Broth (Miller) | BioShop | LBL407.1 |  |
| Tween-80 | BioShop | TWN507.500 | CAS 9005-65-6 |
| Tris base | Sigma | 10708976001 | CAS 77-86-1 |
| Calcium chloride dihydrate | Fisher | BP510-500 | CAS 10035-04-8 |
| Sodium phosphate monobasic monohydrate | BioShop | SPM400.1 | CAS 10049-21-5 |
| Sodium phosphate dibasic heptahydrate | BioShop | SPD579.1 | CAS 7782-85-6 |
| Magnesium sulfate heptahydrate | BioShop | MAG511.1 | CAS 10034-99-8 |
| Lysozyme | BioShop | LYS702.10 | CAS 12650-88-3 |
| Triton X-100 | Sigma | X100-100ML | CAS 9036-19-5 |
| Glycerol | Sigma | G2025 | CAS 56-81-5 |
| Sodium dodecyl sulfate (SDS) | BioShop | SDS001 | CAS 151-21-3 |
| Bromophenol blue | BioShop | BRO777 | CAS 6262-5-28-9 |
| Coomassie Brilliant Blue R-250 | BioShop | CBB250 | CAS 6104-59-2 |
| Citric acid monohydrate | BioShop | CIT003 | CAS 5949-29-1 |
| Ammonium sulfate | BioShop | AMP302 | CAS 7783-20-2 |
| Ellagic acid | Sigma | E2250 | CAS 476-66-4 |
| Urolithin M5 | TRC | TRC-U847035-5MG | CAS 91485-02-8 |
| Urolithin M6 | TRC | U847040 | CAS 1006683-97-1 |
| Urolithin M7 | TRC | TRCU847045-2.5MG | CAS 531512-26-2 |
| Urolithin D | TRC | TRC-U847020-5MG | CAS 131086-98-1 |
| Urolithin C | TRC | U847015 | CAS 165393-06-6 |
| Urolithin A | Sigma | SML1791-5MG | CAS 1143-70-0 |
| Isourolithin A | TRC | I917520 | CAS 174023-48-4 |
| Urolithin B | TRC | U847005 | CAS 1139-83-9 |
| Salicylic acid | Sigma | 84210-100G | CAS 69-72-7 |
| TRI reagent | Zymo Research | R2050-1-50 |  |
| NADH disodium salt, trihydrate, reduced | BioBasic | NB0642.SIZE.1g | CAS 606-68-8 |
| dNTP Mixture (10 mM) | BioBasic | DD0056 |  |
| Ampicillin, sodium salt | BioBasic | AB0028 | CAS 69-52-3 |
| Chloramphenicol | BioBasic | CB0118 | CAS 56-75-7 |
| Tetracycline HCl | BioShop | TET701-10 | CAS 64-75-5 |
| Thiamphenicol | Fisher | J63575.03 | CAS 15318-45-3 |
| Theophylline | Sigma | T1633-50G | CAS 58-55-9 |
| D-cycloserine | Abcam | AB120121 | CAS 68-41-7 |
| Agarose | Fisher | BP160-500 | CAS 9012-36-6 |
| Acrylamide/Bis-acrylamide (29:1) | BioShop | ACR009.500 |  |
| Ammonium persulfate | Sigma | 248614-500G | CAS 7727-54-0 |
| TEMED | BioShop | TEM001.25 | CAS 110-18-9 |
| PBS pH 7.4 (10X) | Gibco | 70011-044 |  |
| Nuclease-free water | Invitrogen | 10977-015 |  |
| MilliQ water system | Millipore | IQ7000 |  |

**Supplementary Table 2.** Bacteria used in this study.

| Name | Strain | Source | Additional Information |
| --- | --- | --- | --- |
| *Clostridium sporogenes* | DSM 795 | DSMZ | Type strain \| Used for heterologous expression |
| *Enterocloster asparagiformis* | DSM 15981 | DSMZ | Type strain |
| *Enterocloster bolteae* | DSM 15670 | DSMZ | Type strain |
| *Enterocloster citroniae* | DSM 19261 | DSMZ | Type strain |
| *Enterocloster pacaense* | CCUG 71489T | CCUG | Type strain – Also published as *Lachnoclostridium pacaense*<sup>67</sup> and *Enterocloster hominis*<sup>68</sup> |
| *Escherichia coli* | NEB10β | NEB | Used for routine cloning and plasmid maintenance |
|  | CA434 | SBRC Nottingham | Used for conjugation with *C. sporogenes* |
| *Gordonibacter pamelaee* | DSM 19378 | DSMZ | Type strain |
| *Gordonibacter urolithinifaciens* | DSM 27213 | DSMZ | Type strain |
| *Gordonibacter* sp. 28C | DSM 110925 | DSMZ |  |
| *Gordonibacter massiliensis* | CSUR P2775 | CSUR | Type strain |
| *Rhodococcus erythropolis* | DSM 43066 | DSMZ | Type strain \| Used for heterologous expression |
| *Rhodococcus erythropolis* pTipQC2-MCS |  | This study | Heterologous expression: multiple cloning site (MCS) |
| *Rhodococcus erythropolis* pTipQC2-Ea_ *ucdCFO* |  | This study | Heterologous expression: *E. asparagiformis ucd* operon proteins |
| *Clostridium sporogenes* pRG-Ea_ *uxdFOC* |  | This study | Heterologous expression: *E. asparagiformis uxd* operon proteins |

DSMZ: German Collection of Microorganisms and Cell Cultures CCUG: Culture Collection University Of Gothenburg NEB: New England Biolabs SBRC: Synthetic Biology Research Centre CSUR: IHU Méditerranée Infection Collection of Microorganisms

**Supplementary Table 3.**
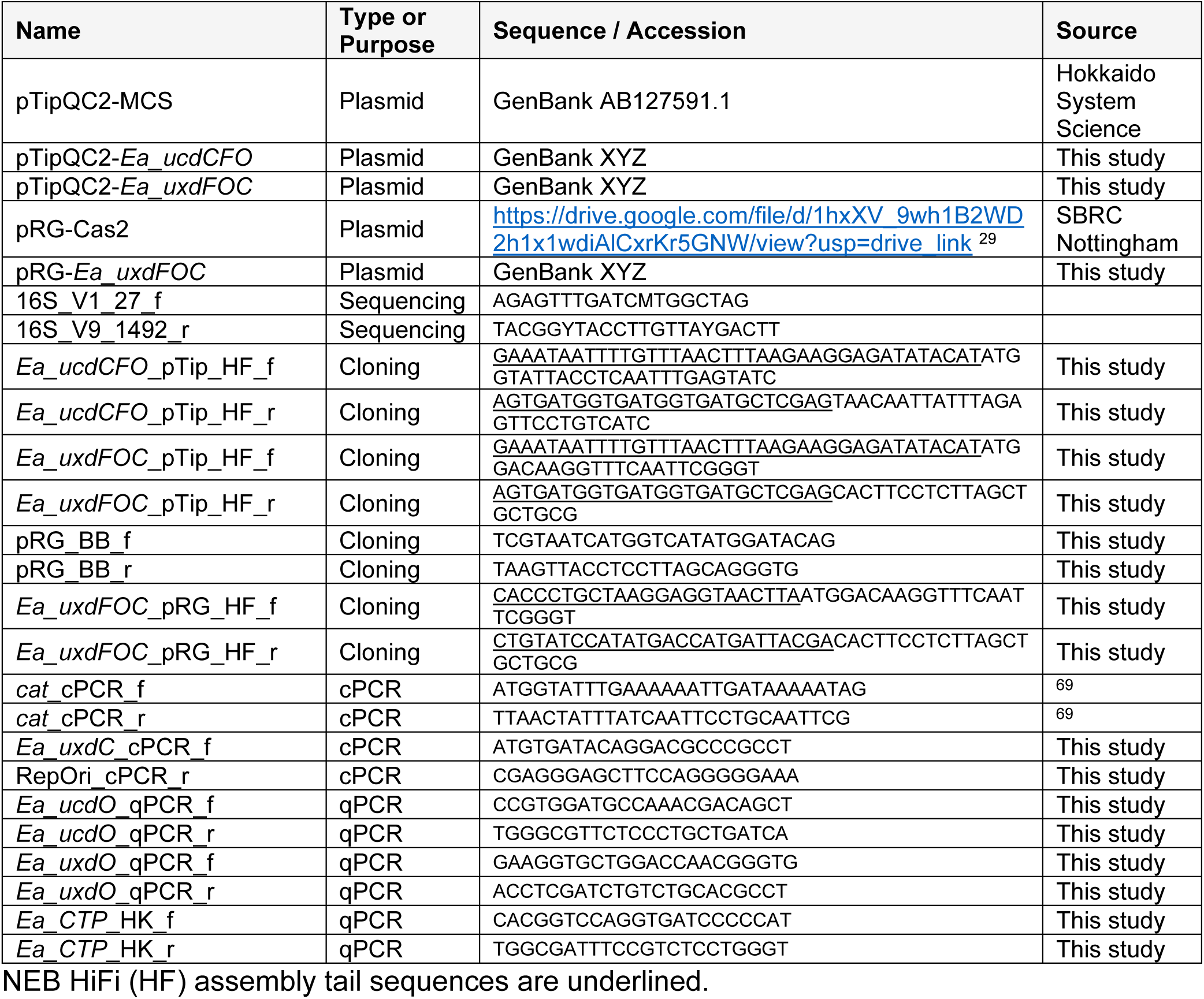
Plasmids and primers used in this study.

**Supplementary Table 4.** PCR conditions for cloning and qPCR.

| **Amplicon** | **Forward** | **Reverse** | **Kit** | **T<sub>a</sub>**<br>**(°C)** | **Extension**<br>**(min:s)** |
| --- | --- | --- | --- | --- | --- |
| *Ea_ucdCFO* pTip_HF | *Ea_ucdCFO* pTip_HF_f | *Ea_ucdCFO* pTip_HF_r | Q5 | 60 | 2:40 |
| *Ea_uxdFOC* pTip_HF | *Ea_uxdFOC* pTip_HF_f | *Ea_uxdFOC* pTip_HF_r | Q5 | 65 | 2:40 |
| pRG_BB | pRG_BB_f | pRG_BB_r | Q5 | 64 | 2:20 |
| *Ea_uxdFOC* pRG_HF | *Ea_uxdFOC* pRG_HF_f | *Ea_uxdFOC* pRG_HF_r | Q5 | 68 | 3:00 |
| *cat*_cPCR | *cat*_cPCR_f | *cat*_cPCR_r | OneTaq<br>2X | 45 | 0:45 |
| *uxdC*-RepOri_cPCR | *Ea_uxdC*_cPCR_f | RepOri_cPCR_r | OneTaq<br>2X | 60 | 1:40 |
| *Ea_ucdO*_qPCR | *Ea_ucdO*_qPCR_f | *Ea_ucdO*_qPCR_r | Luna<br>qPCR | 60 | 0:30 |
| *Ea_uxdO*_qPCR | *Ea_uxdO*_qPCR_f | *Ea_uxdO*_qPCR_r | Luna<br>qPCR | 60 | 0:30 |
| *Ea_CTP*_qPCR<br>(Reference) | *Ea_CTP*_Ref_f | *Ea_CTP*_Ref_r | Luna<br>qPCR | 60 | 0:30 |

**Supplementary Table 5.**
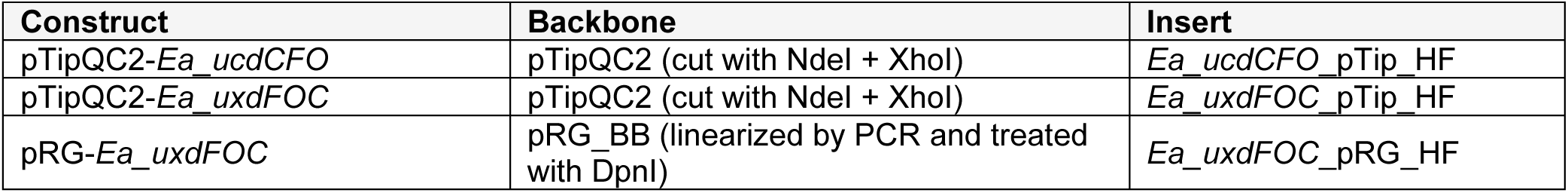
HiFi DNA assembly constructs.

**Supplementary Table 6.** Genome accessions for Fig. 4a.

| **GTDB v10-RS226**<br>**Taxonomy (Species)** | **Strain ID** | **GenBank Assembly** |
| --- | --- | --- |
| *Altitiscatomonas aceti* | BX10 | GCA_014385265.1 |
| *Altitiscatomonas aceti* | P053L1-22 | GCA_037370865.1 |
| *Enterocloster alcoholdehydrogenati* | C5-48 | GCA_013282095.1 |
| *Enterocloster aldenensis* | AF19-9LB | GCA_003434055.1 |
| *Enterocloster aldenensis* | MSK.1.13 | GCA_011317135.1 |
| *Enterocloster aldenensis* | MSK.1.17 | GCA_013304305.1 |
| *Enterocloster aldenensis* | SIG283 | GCA_015152415.1 |
| *Enterocloster aldenensis* | DFI.3.50 | GCA_020554935.1 |
| *Enterocloster aldenensis* | DFI.6.55 | GCA_022136055.1 |
| *Enterocloster aldenensis* | TF09-12AC | GCA_027690515.1 |
| *Enterocloster aldenensis* | ET318 | GCA_030373645.1 |
| *Enterocloster aldenensis* | SUG2707 | GCA_034113485.1 |
| *Enterocloster aldenensis* | S_154 | GCA_034122565.1 |
| *Enterocloster aldenensis* | S_52 | GCA_034124805.1 |
| *Enterocloster aldenensis* | S_22 | GCA_034125605.1 |
| *Enterocloster aldenensis* | S_543 | GCA_034126345.1 |
| *Enterocloster aldenensis* | S_537 | GCA_034126445.1 |
| *Enterocloster aldenensis* | S_347 | GCA_034130145.1 |
| *Enterocloster aldenensis* | S_292 | GCA_034131165.1 |
| *Enterocloster aldenensis* | S_730 | GCA_034133045.1 |
| *Enterocloster aldenensis* | S_694 | GCA_034133765.1 |
| *Enterocloster aldenensis* | S_931 | GCA_034138745.1 |
| *Enterocloster aldenensis* | CM710M_44 | GCA_036318105.1 |
| *Enterocloster aldenensis* | CLA-JM-H53 | GCA_040097765.1 |
| *Enterocloster aldenensis* | RFS30 | GCA_040307795.1 |
| *Enterocloster aldenensis* | 61-73 | GCA_041222985.1 |
| *Enterocloster aldenensis* | extra-ERR1762133.4_1677761238 | GCA_949460435.1 |
| *Enterocloster aldenensis* | extra-ERR1762121.15_1677761188 | GCA_949463055.1 |
| *Enterocloster aldenensis* | extra-ERR1762134.7_1677761246 | GCA_949465065.1 |
| *Enterocloster aldenensis* | extra-ERR1762131.5_1677761229 | GCA_949465575.1 |
| *Enterocloster aldenensis* | extra-ERR1762132.13_1677761236 | GCA_949465705.1 |
| *Enterocloster aldenensis* | extra-ERR1762136.10_1677761251 | GCA_949474955.1 |
| *Enterocloster aldenensis* | extra-ERR1762119.4_1677761174 | GCA_949499535.1 |
| *Enterocloster aldenensis* | extra-ERR1762120.22_1677761183 | GCA_949510155.1 |
| *Enterocloster aldenensis* | ena-SAMPLE-TAB-10-01-2024-09:53:06:039-238938 | GCA_963921755.1 |
| *Enterocloster aldenensis* | CC00468 | GCA_964244085.1 |
| *Enterocloster asparagiformis* | DSM 15981 | GCA_000158075.1 |
| *Enterocloster asparagiformis* | AF04-15 | GCA_003466005.1 |
| *Enterocloster asparagiformis* | DSM 15981 | GCA_025149125.1 |
| *Enterocloster asparagiformis* | S_293 | GCA_034131145.1 |
| *Enterocloster asparagiformis* | ADS30 | GCA_040307595.1 |
| *Enterocloster asparagiformis* | ERR1190745-bin.24 | GCA_905193965.1 |
| *Enterocloster bolteae* | ATCC BAA-613 | GCA_000154365.1 |
| *Enterocloster bolteae* | 90B8 | GCA_000371645.1 |
| *Enterocloster bolteae* | 90B7 | GCA_000371665.1 |
| *Enterocloster bolteae* | 90B3 | GCA_000371685.1 |
| *Enterocloster bolteae* | 90A9 | GCA_000371705.1 |
| *Enterocloster bolteae* | 90A5 | GCA_000371725.1 |
| *Enterocloster bolteae* |  | GCA_000431175.1 |
| *Enterocloster bolteae* | WAL-14578 | GCA_001078425.1 |
| *Enterocloster bolteae* | ATCC BAA-613 | GCA_002234575.2 |
| *Enterocloster bolteae* | ATCC BAA-613 | GCA_002959675.1 |
| *Enterocloster bolteae* | TF09-4AC | GCA_003437595.1 |
| *Enterocloster bolteae* | AF27-9 | GCA_003458165.1 |
| *Enterocloster bolteae* | AF24-13 | GCA_003458625.1 |
| *Enterocloster bolteae* | AF14-18 | GCA_003464745.1 |
| *Enterocloster bolteae* | AM35-14 | GCA_003467855.1 |
| *Enterocloster bolteae* | CBBP-2 | GCA_013112035.1 |
| *Enterocloster bolteae* | 1001302B_160321_D9 | GCA_015547745.1 |
| *Enterocloster bolteae* | D31t1_170403_B11 | GCA_015551555.1 |
| *Enterocloster bolteae* | D40t1_170626_F8 | GCA_015556085.1 |
| *Enterocloster bolteae* | 1001287H_170206_C8 | GCA_015557875.1 |
| *Enterocloster bolteae* | 1001713B170214_170313_G12 | GCA_015558345.1 |
| Enterocloster bolteae | FDAARGOS_1231 | GCA_016889665.1 |
| Enterocloster bolteae | L3_079_062G1_dasL3_079_062G1_metabat.metabat.9 | GCA_018368145.1 |
| Enterocloster bolteae | L3_133_361G1_dasL3_133_361G1_maxbin2.maxbin.016sta_sub | GCA_018373655.1 |
| Enterocloster bolteae | DFI.2.20 | GCA_020554465.1 |
| Enterocloster bolteae | DFI.2.21 | GCA_020555225.1 |
| Enterocloster bolteae | SL.1.21 | GCA_020555685.1 |
| Enterocloster bolteae | DFI.1.195 | GCA_020559295.1 |
| Enterocloster bolteae | DFI.1.207 | GCA_020563055.1 |
| Enterocloster bolteae | AHG0001 | GCA_020736955.1 |
| Enterocloster bolteae | S19_Bin2 | GCA_021771435.1 |
| Enterocloster bolteae | DFI.1.85 | GCA_022137355.1 |
| Enterocloster bolteae | DFI.1.209 | GCA_022138045.1 |
| Enterocloster bolteae | DFI.1.16 | GCA_022138265.1 |
| Enterocloster bolteae | VE303-01 | GCA_023008325.1 |
| Enterocloster bolteae | SL.2.16 | GCA_024460255.1 |
| Enterocloster bolteae | DFI.6.108 | GCA_024462735.1 |
| Enterocloster bolteae | DSM_29485 | GCA_024622735.1 |
| Enterocloster bolteae | AM60-12MH-1A | GCA_027668515.1 |
| Enterocloster bolteae | AM27-28LB | GCA_027671045.1 |
| Enterocloster bolteae | N1154.17 | GCA_030839815.1 |
| Enterocloster bolteae | N1158.18 | GCA_030844385.1 |
| Enterocloster bolteae | N4_188_045G1_dasN4_188_045G1_concoct_22 | GCA_032472135.1 |
| Enterocloster bolteae | N5_242_000G1_dasN5_242_000G1_maxbin2.maxbin.008 | GCA_032501265.1 |
| Enterocloster bolteae | S_214 | GCA_034121425.1 |
| Enterocloster bolteae | S_195 | GCA_034121745.1 |
| Enterocloster bolteae | S_194 | GCA_034121765.1 |
| Enterocloster bolteae | S_185 | GCA_034121915.1 |
| Enterocloster bolteae | S_156 | GCA_034122505.1 |
| Enterocloster bolteae | S_150 | GCA_034122605.1 |
| Enterocloster bolteae | S_87 | GCA_034124105.1 |
| Enterocloster bolteae | S_80 | GCA_034124305.1 |
| Enterocloster bolteae | S_10 | GCA_034125805.1 |
| Enterocloster bolteae | S_4 | GCA_034125985.1 |
| Enterocloster bolteae | S_568 | GCA_034126065.1 |
| Enterocloster bolteae | S_550 | GCA_034126255.1 |
| Enterocloster bolteae | S_488 | GCA_034128065.1 |
| Enterocloster bolteae | S_451 | GCA_034128085.1 |
| Enterocloster bolteae | S_411 | GCA_034128865.1 |
| Enterocloster bolteae | S_393 | GCA_034129245.1 |
| Enterocloster bolteae | S_391 | GCA_034129265.1 |
| Enterocloster bolteae | S_375 | GCA_034129605.1 |
| Enterocloster bolteae | S_365 | GCA_034129785.1 |
| Enterocloster bolteae | S_275 | GCA_034131485.1 |
| Enterocloster bolteae | S_759 | GCA_034132485.1 |
| Enterocloster bolteae | S_752 | GCA_034132625.1 |
| Enterocloster bolteae | S_748 | GCA_034132685.1 |
| Enterocloster bolteae | S_731 | GCA_034133025.1 |
| Enterocloster bolteae | S_679 | GCA_034134065.1 |
| Enterocloster bolteae | S_927 | GCA_034138905.1 |
| Enterocloster bolteae | S_919 | GCA_034139205.1 |
| Enterocloster bolteae | S_903 | GCA_034139785.1 |
| Enterocloster bolteae | S_895 | GCA_034140145.1 |
| Enterocloster bolteae | S_884 | GCA_034140645.1 |
| Enterocloster bolteae | S_859 | GCA_034141645.1 |
| Enterocloster bolteae | S_844 | GCA_034142205.1 |
| Enterocloster bolteae | S_831 | GCA_034142765.1 |
| Enterocloster bolteae | S_822 | GCA_034142925.1 |
| Enterocloster bolteae | S_819 | GCA_034143005.1 |
| Enterocloster bolteae | S_765 | GCA_034144985.1 |
| Enterocloster bolteae | CLA-AA-H15 | GCA_040095555.1 |
| Enterocloster bolteae | RTP31004st1_E3_RTP31004_210507 | GCA_040920825.1 |
| Enterocloster bolteae | MGYG-HGUT-01493 | GCA_902375545.1 |
| Enterocloster bolteae | SRR413704-bin.8 | GCA_905197355.1 |
| Enterocloster bolteae | extra-SRR8581402.5_1677803360 | GCA_949494455.1 |
| Enterocloster bolteae | ERR9353914_bin.7_MetaWRAP_v1.3_MAG | GCA_958411475.1 |
| Enterocloster bolteae | ERR9578304_bin.13_MetaWRAP_v1.3_MAG | GCA_958422245.1 |
| Enterocloster bolteae | ERR9609115_bin.4_MetaWRAP_v1.3_MAG | GCA_958446755.1 |
| Enterocloster bolteae | SRR17798029_bin.1_MetaWRAP_v1.3_MAG | GCA_958451465.1 |
| Enterocloster bolteae | SRR13494490_bin.8_MetaWRAP_v1.3_MAG | GCA_959023705.1 |
| Enterocloster bolteae | ERR10149231 bin.7 MetaWRAP v1.3 MAG | GCA_959025025.1 |
| Enterocloster bolteae | SRR16280003 bin.20 MetaWRAP v1.3 MAG | GCA_959605845.1 |
| Enterocloster bolteae | ERR9969170 bin.30 MetaWRAP v1.3 MAG | GCA_963529055.1 |
| Enterocloster citroniae | WAL-17108 | GCA_000233455.1 |
| Enterocloster citroniae | WAL-19142 | GCA_001078435.1 |
| Enterocloster citroniae | AF29-4 | GCA_003433945.1 |
| Enterocloster citroniae | J1101437_171009_D5 | GCA_015548445.1 |
| Enterocloster citroniae | MCC335 | GCA_018785395.1 |
| Enterocloster citroniae | SL.3.11 | GCA_020554185.1 |
| Enterocloster citroniae | AHG0002 | GCA_020736995.1 |
| Enterocloster citroniae | AH.B8_3 | GCA_021204485.1 |
| Enterocloster citroniae | AM09-31 | GCA_027662505.1 |
| Enterocloster citroniae | AM75-15pH9A | GCA_027666385.1 |
| Enterocloster citroniae | S_157 | GCA_034122445.1 |
| Enterocloster citroniae | CM229S_55 | GCA_036318325.1 |
| Enterocloster citroniae | CLA-AA-H231 | GCA_040094335.1 |
| Enterocloster citroniae | ADS31 | GCA_040306965.1 |
| Enterocloster citroniae | DSM_19261 | GCA_040545455.1 |
| Enterocloster citroniae | RTP31023st1_A12 RTP31023_210422 | GCA_040920865.1 |
| Enterocloster citroniae | RTP21198st1_B4 RTP21198_201120 | GCA_040926685.1 |
| Enterocloster citroniae | NLAE-zl-G70 | GCA_900115855.1 |
| Enterocloster citroniae | MGYG-HGUT-00198 | GCA_902364255.1 |
| Enterocloster citroniae | SRR5713947-bin.24 | GCA_905212655.1 |
| Enterocloster citroniae | ERR4918920 bin.7 MetaWRAP v1.3 MAG | GCA_958443955.1 |
| Enterocloster clostridioformis | 2_1_49FAA | GCA_000234155.1 |
| Enterocloster clostridioformis | CM201 | GCA_000371405.1 |
| Enterocloster clostridioformis | 90B1 | GCA_000371485.1 |
| Enterocloster clostridioformis | 90A8 | GCA_000371505.1 |
| Enterocloster clostridioformis | 90A6 | GCA_000371545.1 |
| Enterocloster clostridioformis | 90A4 | GCA_000371565.1 |
| Enterocloster clostridioformis | 90A3 | GCA_000371585.1 |
| Enterocloster clostridioformis | 90A1 | GCA_000371605.1 |
| Enterocloster clostridioformis |  | GCA_000436455.1 |
| Enterocloster clostridioformis | WAL-7855 | GCA_001078445.1 |
| Enterocloster clostridioformis | 2789STDY5834865 | GCA_001405335.1 |
| Enterocloster clostridioformis | 1001175st1_C5 | GCA_005844705.1 |
| Enterocloster clostridioformis | NBRC_113352 | GCA_006538465.1 |
| Enterocloster clostridioformis | FDAARGOS_739 | GCA_012273195.1 |
| Enterocloster clostridioformis | MSK.2.80 | GCA_013299965.1 |
| Enterocloster clostridioformis | MSK.2.98 | GCA_013304125.1 |
| Enterocloster clostridioformis | MSK.2.94 | GCA_013304155.1 |
| Enterocloster clostridioformis | MSK.2.90 | GCA_013304185.1 |
| Enterocloster clostridioformis | MSK.2.63 | GCA_013304205.1 |
| Enterocloster clostridioformis | MSK.2.78 | GCA_013304225.1 |
| Enterocloster clostridioformis | MSK.2.73 | GCA_013304245.1 |
| Enterocloster clostridioformis | MSK.2.59 | GCA_013304255.1 |
| Enterocloster clostridioformis | MSK.2.26 | GCA_013304275.1 |
| Enterocloster clostridioformis | SIG284 | GCA_015152355.1 |
| Enterocloster clostridioformis | 1001175B_160314_C5 | GCA_015548065.1 |
| Enterocloster clostridioformis | J1101653_170612_A4 | GCA_015669475.1 |
| Enterocloster clostridioformis | L2_031_030G1_dasL2_031_030G1_metabat.metabat.6 | GCA_018381395.1 |
| Enterocloster clostridioformis | CD33_MAG32 | GCA_019012735.1 |
| Enterocloster clostridioformis | FDAARGOS_1529 | GCA_020297485.1 |
| Enterocloster clostridioformis | ET46 | GCA_021532095.1 |
| Enterocloster clostridioformis | SUG543 | GCA_022771365.1 |
| Enterocloster clostridioformis | AFF13-24A | GCA_027663445.1 |
| Enterocloster clostridioformis | D35st1_B1_D35t1_190705 | GCA_028210245.1 |
| Enterocloster clostridioformis | D35st1_B2_D35t1_190705 | GCA_028210555.1 |
| Enterocloster clostridioformis | J1101653st1_B2_J1101653_170612 | GCA_028210575.1 |
| Enterocloster clostridioformis | J1100826st1_G3_J1100826_190719 | GCA_028210655.1 |
| Enterocloster clostridioformis | N4_139_000G1_dasN4_139_000G1_concoct_21 | GCA_032477495.1 |
| Enterocloster clostridioformis | SUG2708 | GCA_034115905.1 |
| Enterocloster clostridioformis | S_279 | GCA_034131425.1 |
| Enterocloster clostridioformis | S_592 | GCA_034136325.1 |
| Enterocloster clostridioformis | P045L1-6 | GCA_037353045.1 |
| Enterocloster clostridioformis | RTP21489st1_B4 RTP31083_211112 | GCA_039753315.1 |
| Enterocloster clostridioformis | RTP21489st1_A4 RTP21489_200313 | GCA_039753355.1 |
| Enterocloster clostridioformis | J1101653st1_H5_J1100826_190719 | GCA_039837225.1 |
| Enterocloster clostridioformis | J1101653st1_F9_J1101653_170612 | GCA_039837275.1 |
| Enterocloster clostridioformis | J1100826st1_D9_J1100826_190719 | GCA_039837315.1 |
| Enterocloster clostridioformis | CLA-JM-H51 | GCA_040097865.1 |
| Enterocloster clostridioformis | RFS31 | GCA_040307325.1 |
| Enterocloster clostridioformis | RTP31106st1_B11_RTP31106_210429 | GCA_040915925.1 |
| Enterocloster clostridioformis | RTP31106st1_C9_RTP31106_210429 | GCA_040915945.1 |
| Enterocloster clostridioformis | RTP31106st1_G10_RTP31106_210429 | GCA_040915965.1 |
| Enterocloster clostridioformis | RTP31106st1_A6_RTP31106_210429 | GCA_040916005.1 |
| Enterocloster clostridioformis | RTP31106st1_A10_RTP31106_210429 | GCA_040916045.1 |
| Enterocloster clostridioformis | D5252bin20 | GCA_041327215.1 |
| Enterocloster clostridioformis | NLAE-zl-C196 | GCA_900100685.1 |
| Enterocloster clostridioformis | NLAE-zl-G208 | GCA_900108895.1 |
| Enterocloster clostridioformis | ATCC 25537 | GCA_900113155.1 |
| Enterocloster clostridioformis | NCTC11224 | GCA_900447015.1 |
| Enterocloster clostridioformis | MGYG-HGUT-01386 | GCA_902374585.1 |
| Enterocloster clostridioformis | MGBC136599 | GCA_910586265.1 |
| Enterocloster clostridioformis | MGBC104247 | GCA_948475675.1 |
| Enterocloster clostridioformis | MGBC103138 | GCA_948511265.1 |
| Enterocloster clostridioformis | MGBC103121 | GCA_948522125.1 |
| Enterocloster clostridioformis | MGBC103132 | GCA_948524135.1 |
| Enterocloster clostridioformis | MGBC104273 | GCA_948582345.1 |
| Enterocloster clostridioformis | MGBC103192 | GCA_948587995.1 |
| Enterocloster clostridioformis | MGBC103136 | GCA_948603615.1 |
| Enterocloster clostridioformis | MGBC136614 | GCA_948661395.1 |
| Enterocloster clostridioformis | MGBC104354 | GCA_948681025.1 |
| Enterocloster clostridioformis | MGBC102828 | GCA_948901585.1 |
| Enterocloster clostridioformis | MGBC112024 | GCA_948908805.1 |
| Enterocloster clostridioformis | MGBC102824 | GCA_948919595.1 |
| Enterocloster clostridioformis | MGBC102818 | GCA_948932345.1 |
| Enterocloster clostridioformis | MGBC140751 | GCA_948952455.1 |
| Enterocloster clostridioformis | MGBC140627 | GCA_949013715.1 |
| Enterocloster clostridioformis | MGBC112249 | GCA_949066195.1 |
| Enterocloster clostridioformis | MGBC112317 | GCA_949081035.1 |
| Enterocloster clostridioformis | MGBC112283 | GCA_949121295.1 |
| Enterocloster clostridioformis | NCI_12_NClco.18_1677235885 | GCA_949343865.1 |
| Enterocloster clostridioformis | extra-SRR6945388.13_1677799424 | GCA_949390495.1 |
| Enterocloster clostridioformis | extra-SRR6945379.5_1677799372 | GCA_949400645.1 |
| Enterocloster clostridioformis | extra-SRR6945382.10_1677799393 | GCA_949474175.1 |
| Enterocloster clostridioformis | extra-SRR6945416.15_1677800340 | GCA_949523935.1 |
| Enterocloster clostridioformis | extra-SRR6945408.8_1677800313 | GCA_949531885.1 |
| Enterocloster clostridioformis | extra-SRR6945412.8_1677800327 | GCA_949542115.1 |
| Enterocloster clostridioformis | iMGMC-40_1678881685 | GCA_949742355.1 |
| Enterocloster clostridioformis | ERR9762695_bin.2_MetaWRAP_v1.3_MAG | GCA_958414435.1 |
| Enterocloster clostridioformis | SRR18940293_bin.38_MetaWRAP_v1.3_MAG | GCA_958435425.1 |
| Enterocloster clostridioformis | SRR17798039_bin.8_MetaWRAP_v1.3_MAG | GCA_958436665.1 |
| Enterocloster clostridioformis | SRR22541663_bin.17_MetaWRAP_v1.3_MAG | GCA_958453805.1 |
| Enterocloster clostridioformis | ena-SAMPLE-TAB-10-01-2024-09:53:06:039-238937 | GCA_963921695.1 |
| Enterocloster clostridioformis_A | AGR2157 | GCA_000424325.1 |
| Enterocloster clostridioformis_A | ERR10149264_bin.22_MetaWRAP_v1.3_MAG | GCA_959023645.1 |
| Enterocloster excrementigallinarum | CHK198-12963 | GCA_019119955.1 |
| Enterocloster excrementigallinarum | BRZ_GD_bin72 | GCA_944382075.1 |
| Enterocloster excrementigallinarum | dsm_us_coassembly_bin138 | GCA_947854055.1 |
| Enterocloster excrementipullorum | CHK180-15479 | GCA_019119575.1 |
| Enterocloster excrementipullorum | BRZ_CH_bin70 | GCA_944384005.1 |
| Enterocloster excrementipullorum | metabat2_XCR1_HOM_7852_bwa.48_MAG | GCA_963928035.1 |
| Enterocloster faecavium | CHK188-4685 | GCA_019118585.1 |
| Enterocloster faecavium | BRZ_IC_bin19 | GCA_944381665.1 |
| Enterocloster lavalensis | KCTC 15153 | GCA_003024655.1 |
| Enterocloster lavalensis | SL.2.12 | GCA_020564305.1 |
| Enterocloster lavalensis | D6_SB_15 | GCA_031293285.1 |
| Enterocloster lavalensis | S_54 | GCA_034124785.1 |
| Enterocloster lavalensis | CTOTU13361 | GCA_036750505.1 |
| Enterocloster lavalensis | RTP21230st1_H1_RTP21230_210325 | GCA_040926545.1 |
| Enterocloster lavalensis | RTP21230st1_F6_RTP21230_210325 | GCA_040926585.1 |
| Enterocloster lavalensis | RTP21230st1_A7_RTP21230_210325 | GCA_040926665.1 |
| Enterocloster lavalensis | NLAE-zl-G277 | GCA_900102595.1 |
| Enterocloster lavalensis | MGYG-HGUT-00172 | GCA_902364025.1 |
| Enterocloster lavalensis | ERR1190731-bin.3 | GCA_905194475.1 |
| Enterocloster lavalensis | MTG234_bin.53.fa | GCA_934852855.1 |
| Enterocloster lavalensis | MTG246_bin.51.fa | GCA_934882645.1 |
| Enterocloster lavalensis | MTG236_bin.16.fa | GCA_934882855.1 |
| Enterocloster sp000155435 | DFI.6.51A | GCA_020709355.1 |
| Enterocloster sp000155435 | SL.2.13 | GCA_020709765.1 |
| Enterocloster sp000155435 | OA13 | GCA_022440595.1 |
| Enterocloster sp000155435 | CLA-SR-H021 | GCA_040096035.1 |
| Enterocloster sp000431375 | Gw_UH_bin_249 | GCA_018053045.1 |
| Enterocloster sp000431375 | DOME_MAG_12367 | GCA_036301185.1 |
| Enterocloster sp000431375 | P002L1-10 | GCA_037234785.1 |
| Enterocloster sp000431375 | P015-4 | GCA_037247195.1 |
| Enterocloster sp000431375 | P022L1-36 | GCA_037277075.1 |
| Enterocloster sp000431375 | P027-2 | GCA_037295135.1 |
| Enterocloster sp000431375 | P029-2 | GCA_037302955.1 |
| Enterocloster sp000431375 | P028L1-22 | GCA_037304265.1 |
| Enterocloster sp000431375 | P032L1-17 | GCA_037317655.1 |
| Enterocloster sp000431375 | P030-4 | GCA_037322975.1 |
| Enterocloster sp000431375 | P042L1-25 | GCA_037337525.1 |
| Enterocloster sp000431375 | P037L1-19 | GCA_037348905.1 |
| Enterocloster sp000431375 | P050-24 | GCA_037358175.1 |
| Enterocloster sp000431375 | P047L1-40 | GCA_037366705.1 |
| Enterocloster sp000431375 | P053L1-49 | GCA_037370285.1 |
| Enterocloster sp000431375 | P058L1-12 | GCA_037426565.1 |
| Enterocloster sp000431375 | D1217bin36 | GCA_041323175.1 |
| Enterocloster sp000431375 | D1151bin22 | GCA_041324515.1 |
| Enterocloster sp000431375 | D1125bin31 | GCA_041325895.1 |
| Enterocloster sp000431375 | D1020bin14 | GCA_041327035.1 |
| Enterocloster sp000431375 | D5214bin15 | GCA_041327705.1 |
| Enterocloster sp000431375 | D3096bin9 | GCA_041331655.1 |
| Enterocloster sp000431375 | D3029bin66 | GCA_041333045.1 |
| Enterocloster sp000431375 | D3002bin21 | GCA_041333915.1 |
| Enterocloster sp000431375 | D3037bin7 | GCA_041335475.1 |
| Enterocloster sp000431375 | REFINED_METABAT215_SUBJECT_CONTIGS_1500_ASSEMBLY_K77_MERGED_Pilot_MoBio_Fiber_N_24_7019.54 | GCA_934358685.1 |
| Enterocloster sp000431375 | REFINED_METABAT215_TOP10_CONTIGS_1500_ASSEMBLY_K77_MERGED_Nepal_MoBio_Fiber-Hadza-Nepal_H_23_RAJ1014YZ.78 | GCA_934376865.1 |
| Enterocloster sp000431375 | REFINED_METABAT215_TOP10_CONTIGS_1500_ASSEMBLY_K77_MERGED_Hadza_PheChl_Fiber-Hadza-Nepal_E_16_1654.235 | GCA_934437995.1 |
| Enterocloster sp001517625 | Gw_UH_bin_231 | GCA_018055585.1 |
| Enterocloster sp001517625 | P014L1-23 | GCA_037247845.1 |
| Enterocloster sp001517625 | P013L1-12 | GCA_037249885.1 |
| Enterocloster sp001517625 | P026L1-6 | GCA_037295595.1 |
| Enterocloster sp001517625 | P032L1-15 | GCA_037317675.1 |
| Enterocloster sp001517625 | P007L1-3 | GCA_037454835.1 |
| Enterocloster sp001517625 | HCN-30185 | GCA_037890305.1 |
| Enterocloster sp001517625 | REFINED_METABAT215_TOP10_CONTIGS_1500_ASSEMBLY_K77_MERGED_Nepal_MoBio_Fiber-Hadza-Nepal_H_21_RAJ1012YZ.28 | GCA_934402005.1 |
| Enterocloster sp001517625 | SRS10476999_bin15 | GCA_963557545.1 |
| Enterocloster sp005845215 | OA11 | GCA_022440645.1 |
| Enterocloster sp900541315 | DOME_MAG_10320 | GCA_036305165.1 |
| Enterocloster sp900541315 | bin.509 | GCA_037068625.1 |
| Enterocloster sp900541315 | P018L1-30 | GCA_037256905.1 |
| Enterocloster sp900541315 | P026L1-7 | GCA_037295555.1 |
| Enterocloster sp900541315 | P031L1-7 | GCA_037319585.1 |
| Enterocloster sp900541315 | REFINED_METABAT215_TOP10_CONTIGS_1500_ASSEMBLY_K77_MERGED_Hadza_MoBio_F_H_7_1932.23 | GCA_934387265.1 |
| Enterocloster sp900541315 | REFINED_METABAT215_TOP10_CONTIGS_1500_ASSEMBLY_K77_MERGED_Hadza_PheChl_Fiber-Hadza-Nepal_A_24_1660.405 | GCA_934394355.1 |
| Enterocloster sp900541315 | REFINED_METABAT215_TOP10_CONTIGS_1500_ASSEMBLY_K77_MERGED_Nepal_MoBio_Fiber-Hadza-Nepal_B_3_THA0078JZ.1 | GCA_934408575.1 |
| Enterocloster sp900555905 | L2_037_371G1_dasL2_037_371G1_concoct_55_sub | GCA_018380885.1 |
| Enterocloster sp900555905 | RAGGC_281 | GCA_030824975.1 |
| Enterocloster sp900759225 | ERS329807_bin46 | GCA_963556915.1 |
| Enterocloster sp902464595 | SRS10476986_bin57 | GCA_963560105.1 |
| Enterocloster sp944376135 | E0_LS_bin20 | GCA_944376135.1 |
| Enterocloster sp944382045 | BRZ_LM_bin53 | GCA_944382045.1 |
| Enterocloster sp944383785 | BRZ_PI_bin16 | GCA_944383785.1 |
| Enterocloster sp944384065 | BRZ_CR_bin7 | GCA_944384065.1 |

The full genome accessions table with results for **Fig. 4a** is available in Zenodo [https://doi.org/10.5281/zenodo.16740616].

**Supplementary Table 7.** Genome accessions for Fig. 4b.

| Name in Fig. 4b | Strain ID | GenBank Assembly | GTDB v10-RS226 Taxonomy (Species) |
| --- | --- | --- | --- |
| *Ea* | DSM 15981 | GCA_025149125.1 | *Enterocloster asparagiformis* |
| *Ec* | WAL-19142 | GCA_001078435.1 | *Enterocloster citroniae* |
| *Ep* | CCUG 71489T | GCA_900566185.1 | Enterocloster sp000155435 (previously *Enterocloster pacaense*). Note that strain CCUG 71489T failed quality check in the GTDB. |
| *Eb* | DSM 15670 | GCA_002234575.2 | *Enterocloster bolteae* |

**Supplementary Table 8.** Protein sequence accessions.

| Protein | Species | Mass (Da) | #AA | GenBank | UniProt |
| --- | --- | --- | --- | --- | --- |
| Ea_UcdO | *E. asparagiformis* | 86,310 | 787 | WP_007711966.1 | C0D162 or A0A413FKI6 |
| Eb_UcdO | *E. bolteae* | 86,556 | 787 | WP_002569573.1 | A8RZR2 or A0A414AWH1 or R0B269 |
| Ec_UcdO | *E. citroniae* | 86,383 | 787 | WP_007860285.1 | A0A0J9C6D9 or A0A3E2VBJ6 or G5HFF2 |
| Ep_UcdO | *E. pacaense* | 86,438 | 787 | WP_025486543.1 | NA |
| Ea_UxdO | *E. asparagiformis* | 83,311 | 758 | WP_117776983.1 | A0A413FKH9 |
| Eb_UxdO | *E. bolteae* | 84,349 | 773 | WP_002565238.1 | A8RZP2 |
| Ec_UxdO | *E. citroniae* | 83,243 | 758 | WP_048929765.1 | A0A0J9C6D6 |
| Ep_UxdO | *E. pacaense* | 82,984 | 758 | WP_130789890.1 | NA |
| Ea_UcdC | *E. asparagiformis* | 30,710 | 288 | WP_024734846.1 | A0A413FKP9 |
| Ea_UcdF | *E. asparagiformis* | 17,765 | 163 | WP_007711967.1 | C0D163 or A0A413FKG8 |
| Ea_UxdC | *E. asparagiformis* | 86,960 | 810 | WP_117776984.1 | A0A413FKH3 |
| Ea_UxdF | *E. asparagiformis* | 18,176 | 171 | WP_007711960.1 | C0D156 or A0A413FKR5 |

